# *In situ* cryo-electron tomography reveals the asymmetric architecture of mammalian sperm axonemes

**DOI:** 10.1101/2022.05.15.492011

**Authors:** Zhen Chen, Garrett A. Greenan, Momoko Shiozaki, Yanxin Liu, Will M. Skinner, Xiaowei Zhao, Shumei Zhao, Rui Yan, Caiying Guo, Zhiheng Yu, Polina V. Lishko, David A. Agard, Ronald D. Vale

## Abstract

The flagella of mammalian sperm display non-planar, asymmetric beating, in contrast to the planar, symmetric beating of flagella from sea urchin sperm and unicellular organisms. The molecular basis of this difference is unclear. Here, we perform *in situ* cryo-electron tomography of mouse and human sperm axonemes, providing the highest resolution structural information to date. Our subtomogram averages reveal mammalian sperm-specific protein complexes within the outer microtubule doublets, the radial spokes and nexin-dynein regulatory complexes. The locations and structures of these complexes suggest potential roles in enhancing the mechanical strength of mammalian sperm axonemes and regulating dynein-based axonemal bending. Intriguingly, we find that each of the nine outer microtubule doublets is decorated with a distinct combination of sperm-specific complexes. We propose that this asymmetric distribution of proteins differentially regulates the sliding of each microtubule doublet and may underlie the asymmetric beating of mammalian sperm.

## Introduction

Eukaryotic flagellum and motile cilium are whip-like organelles whose rhythmic beating propels single-cell eukaryotes through fluids, clear dust particles in respiratory tracts, and enable the swimming of sperm cells of various species^1–3^. Most flagella from protozoa to mammals share a conserved core structure, the axoneme, comprised of nine doublet microtubules (doublets) arranged in a circle around a central pair complex of two singlet microtubules (the 9+2 configuration, Fig. 1a)^4, 5^. Dyneins, microtubule-based molecular motors anchored on the nine doublets, drive the relative sliding of neighboring doublets. However, if all dyneins were active at once, forces around the circle of nine outer doublets would be canceled and the structure would not bend^5, 6^. To produce rhythmic beating motions, non-motor protein complexes are needed to regulate dynein activities across the axoneme structure^5, 7–13^. The largest and most critical of these regulatory complexes are the radial spokes that bridge the outer doublets to the central pair complex, and the nexin-dynein regulatory complexes (N-DRCs) that crosslink neighboring doublets and regulate dynein activities across the axoneme structure.

**Figure 1.**
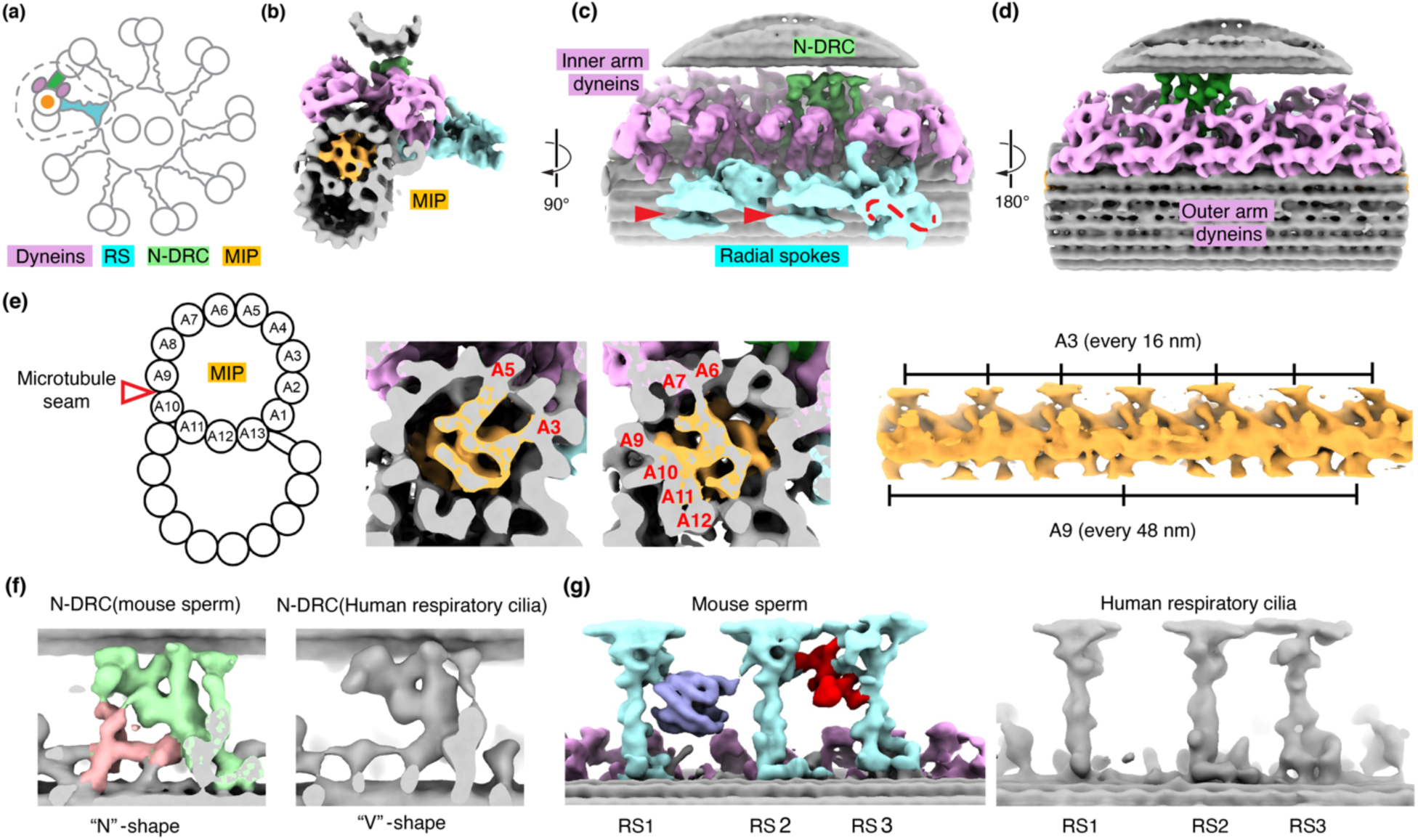
Consensus subvolume averages of doublet microtubules in mouse sperm reveal unique features in non-motor protein complexes including microtubule inner proteins (MIPs), nexin-dynein regulatory complex (N-DRC) and radial spokes (RS). **a,** Schematic of a cross-section view of the conserved (9+2)-microtubule configuration of axonemes in motile cilia. One doublet is highlighted, and its associated motor and non- motor protein complexes are shown. The dyneins, N-DRC, MIPs and RS are colored in pink, green, orange and cyan, respectively. **b-d,** Three views of the global subvolume average of doublet microtubules in mouse sperm axonemes. Different protein complexes are color-coded as in a. The clefts in the top of radial spokes 1 and 2 and the “S”-shaped head of radial spoke 3 are indicated by the red arrowheads and dashed line, respectively in c. **e,** Schematic of the doublet microtubule with individual protofilaments labeled. Two cross-sections of the A-tubule and one longitudinal view of the isolated MIPs densities are shown. The MIPs are colored in orange. The protofilaments connecting to the MIPs and the periodicities for the connections are labeled. **f, g,** Comparison of densities of N- DRC and radial spokes from mouse sperm flagella (this study) and human respiratory cilia (EMDB: 5950) (REF). In f, Additional densities in the mouse sperm N-DRC are highlighted (light red). In g, the unique densities of a barrel and a RS2-RS3 crosslinker in the mouse sperm axoneme are highlighted in blue and red, respectively.

Flagella from different cells display a wide variety of beating patterns, from the planar and symmetric waveforms observed in flagella of unicellular organisms and sea urchin sperm, to the varied non-planar and asymmetric waveforms displayed by different mammalian sperm^6^. The structural and regulatory mechanisms underlying these different waveforms are poorly understood. Much of our current structural understanding of axonemes is derived from studies of *Chlamydomonas* and sea urchin sperm flagellum using advanced cryo-electron tomography (cryo-ET) and image processing^14, 15^. Apart from minor variations of the dyneins on a subset of the nine doublets, most of the other motor and non-motor protein complexes were found to be the same across the nine outer doublets. A unique bridge-like structure that crosslinks two neighboring doublets is proposed to constrain the plane of bending^15, 16^. The pseudo-ninefold symmetry and the bridge structure are thought to be important for generating equivalent beating amplitudes in the opposite directions, leading to planar and symmetric waveforms.

Comparable structural information for mammalian sperm, which display varied non-planar asymmetric waveforms^17–19^, has lagged behind. A technical challenge in using modern cryo-electron microscopy to investigate mammalian sperm flagella is their thickness (>500 nm), which is close to the upper limit for imaging on the widely used 300 kV electron microscopes. Recently, cryogenic focused ion beam/scanning electron microscopy (cryo FIB-SEM) and cryo-ET have been applied to study *in situ* macromolecular structures in sperm axoneme from mammals^20^. However, the limited data obtained in the previous study precluded processing strategies to analyze individual microtubule structures within the axonemes and also their spatial relationships *in situ*.

Here, we combined cryo FIB-SEM and *in situ* cryo-ET with data processing strategies to study the contextual assembly of different microtubule-based structures within mouse and human sperm flagella. Our data provide the highest resolution information to date (18 Å) for the mammalian sperm axonemes, validating the potential of our cellular tomography workflow. Our data reveal non-motor protein complexes in mammalian sperm cells that are not found in axonemes of other mammalian tissues and non-mammalian sperm. We show that each of the nine outer doublets is unique with regard to the composition of regulatory complexes including radial spokes and N-DRC. We propose that the asymmetric distribution of these regulatory complexes across the axoneme could contribute to the asymmetric and nonplanar beating waveforms of various mammalian sperm.

## Results

### Sperm-specific features revealed by subvolume averaging

Freshly extracted mouse sperm were vitrified on EM grids and loaded into a cryo FIB-SEM to generate lamellae of ∼ 300 nm-thickness (Extended Data Fig. 1a, b). The lamellae were then imaged using a Krios 300 kV cryo transmission electron microscope and dose symmetric tomographic tilt series (± 48°) around the axoneme axis were then acquired (N = 108) (Extended Data Fig. 1c). Our images showed detailed molecular features including the double-bilayer membranes of the surrounding mitochondria and individual microtubule protofilaments (Extended Data Fig. 1e,f). 3D tomograms were reconstructed from the tilt series, which revealed repetitive axonemal dyneins and radial spokes along the outer doublets, as well as periodic protrusions from the singlet microtubules of the central pair complex (Extended Data Fig. 1h,i). The periodicities of the radial spokes and central pair protrusions were ∼96 nm and ∼32 nm, respectively (Extended Data Fig. 1i), consistent with those described in *Chlamydomonas* and sea urchin sperm^21, 22^.

To overcome the low signal-noise ratio of raw tomograms, we used subvolume alignment and averaging of repeating units to reconstruct consensus density maps (Fig. 1). Our consensus maps of periodic units from all nine doublets revealed robust signals for individual microtubule protofilaments and other associated protein complexes that repeat every 96 nm (Fig. 1b-d, 24 Å at FSC = 0.143, N = 9099 subvolumes).

Inside the A-tubule of the outer doublet, we observed a filament-like density of microtubule inner proteins (MIPs) that is very similar to, but more extensive than that recently assigned as tektin filaments in bovine trachea cilia^23^ (Fig. 1b and Extended Data Fig. 2). The densities of MIPs have a periodicity of 48 nm, consistent with previously studied *Chlamydomonas* flagella and bovine trachea cilia (18 Å at FSC = 0.143, N = 18153 subvolumes). The filamentous components extend protrusions that connect to all thirteen protofilaments of the A-tubule in mouse sperm axonemes (Extended Data Fig. 2a), in contrast to the more limited connections of tektin filaments to A9-A13 and A1 protofilaments observed in bovine trachea cilia (Fig. 1e and see comparisons in Extended Data Fig. 2b). We observed three different modes of interaction between the MIPs and the lumen of microtubules: 1) interaction with tubulins within a single protofilament, 2) connections to the inter-protofilament space across two neighboring protofilaments, and 3) connections spanning multiple protofilaments (Figure 1e and Extended Data Fig. 2). Notably, the A9-A10 junction is where the microtubule seam of the A-tubule is located^24^, and we observed several sperm-specific densities spanning protofilaments A9-A10 that are absent in the map of bovine respiratory cilia (Extended Data Fig. 2b). Together, our average revealed sperm-specific MIPs that form an extensive network inside the A-tubule of doublet microtubules.

Our consensus map of mouse sperm axoneme reveals similar outer and inner arm dynein structures to those observed in sea urchin sperm (Fig. 1b-d and Extended Data Fig. 3). However, we also observed several unique non-motor protein complexes in the mouse sperm axoneme that do not have an equivalent counterpart in the published axoneme maps from *Chlamydomonas*, *Tetrahymena* and human respiratory cilia (Figure 1f,g)^14, 21, 25, 26^. While the N-DRC in human respiratory cilia has a “V” shape^26^, our consensus map reveals extra densities that extend to the microtubule surface, creating an “N”-shaped structure (Fig. 1f). The radial spokes are comprised of three tower-like densities with two adopting similar morphology (RS1 and RS2) and a third, distinct radial spoke 3 (RS3) (Fig. 1b,c,g). When viewed from the “head” of the towers, radial spoke 1 (RS1) and radial spoke 2 (RS2) both exhibit a gap and a C2 symmetry axis that extends through the “tower”, while the radial spoke 3 (RS3) has a distinctive S-shaped surface with two holes (Fig. 1c). These radial spoke features are similar to the ones observed in human respiratory cilia but differ from the flat radial spoke heads from *Chlamydomonas* and *Tetrahymena*. Multiple additional densities, not found in respiratory cilia or unicellular organisms, were observed between the three spokes in the mouse sperm axoneme (Fig. 1g). First, we observed an ∼20 nm-sized barrel-shaped density between RS1 and RS2, consistent with extra densities in sperm axonemes reported recently^20, 27^. Our higher resolution map revealed that the barrel is comprised of ten rod-shaped strands arranged in a right-handed twist configuration (Supplementary Movie 1). Furthermore, densities were found to crosslink RS2 and RS3, hereafter named as “RS2-RS3 crosslinker” (Fig. 1g). These additional densities could potentially couple the three radial spokes together, which could serve to increase mechanical stability or facilitate coordinated movements. Of note, all of these extra densities in the MIPs, N-DRC and radial spokes are apparent even when our maps were low-pass filtered to 50 Å, a resolution lower than the published map of human respiratory cilia axoneme that does not possess these features (Extended Data Fig. 4). Therefore, the different densities found between respiratory cilia and sperm are not due to higher resolution in this work but most likely reflect the presence of additional sperm-specific proteins.

### The outer doublets are arranged in fixed radial positions around the central pair complex

We next calculated a subvolume average for the 32 nm-repeating units of the central pair complex (Fig.2a-c, 26 Å at FSC = 0.143, N = 3061 subvolumes). Individual protofilaments of the two singlet microtubules were clearly resolved (Fig. 2b). Various proteins protrude from both microtubules, giving rise to a highly asymmetric cross-section contour of the central pair complex (Fig. 2b), which is similar to the central pair complexes from sea urchin sperm^22^. We observed two distinct MIPs inside the two singlet microtubules, both of which repeat every 32 nm (Extended Data Fig. 5a). On the external surface of microtubules, we observed both 32 nm-repeating and 16 nm-repeating features. Notably, compared to the density map for the central pair of sea urchin sperm, where MIPs were not observed, the overall shapes of the external protrusions are very similar, which suggests that the asymmetric surface of the central pair complex is largely conserved from invertebrate to vertebrate sperm (Extended Data Fig. 5b).

**Figure 2.**
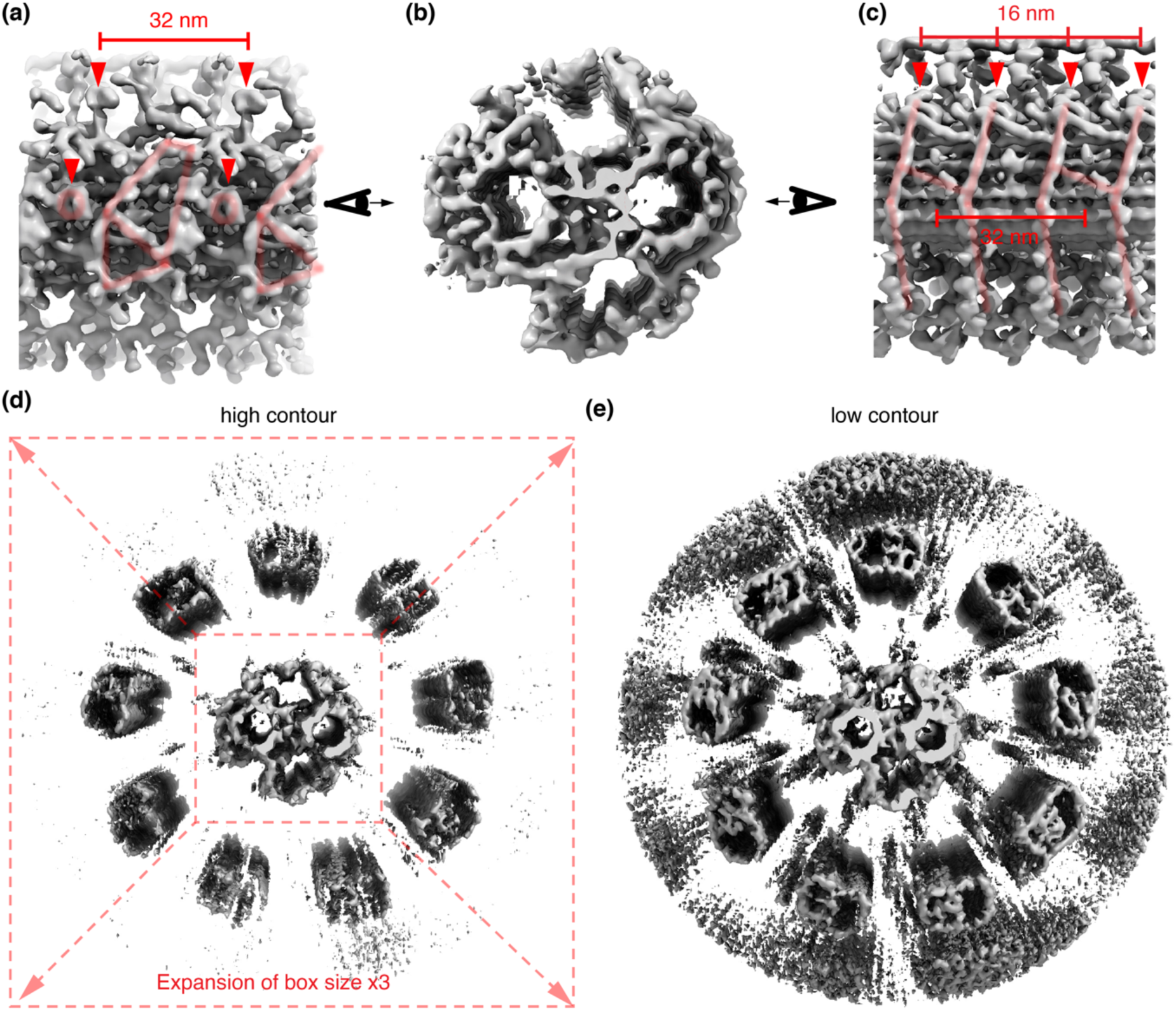
The central pair complex presents asymmetric surfaces in different directions. **a-c**, Three views of the central pair complex of the mouse sperm axoneme. In **a**, **c**, repeating densities are indicated by red arrowheads or highlighted by light red lines and their periodicities are labeled. **d**, An average of the entire axoneme calculated by expanding the original subvomes for the central pair complex three-fold, and averaging without further alignment. At high contour, densities of nine doublet microtubules are resolved at nine distinct radial positions. **e**, The same average as in **d** is shown at low contour. Note the A-tubules from doublet microtubules are distinguishable due to the presence of extensive MIPs densities. Smeared densities corresponding to where dyneins, radial spokes and N-DRCs locate could be observed at lower occupancies than the doublet microtubules.

To understand how the outer doublets are arranged relative to the asymmetric central pair complex, we expanded our aligned subvolumes of the central pair complex three times to include the region where the outer doublet microtubules reside and then calculated an average without further alignment (Fig. 2d,e). Although subvolumes were only aligned for the center central pair complex, nine distinct doublet microtubule densities that are parallel to the singlet microtubules could be resolved, indicating a remarkably consistent radial arrangement of doublets in the axonemes. The A- and B-tubules of doublet microtubule were clearly distinguishable based on the extensive MIP network in the former, but periodic external attachments such as dyneins, radial spokes and N-DRC were poorly resolved, suggesting the lack of longitudinal alignment as might be expected for structures that slide relative to one another (Fig. 2d,e). These analyses of the average of a segment of the entire (9+2) axoneme revealed that the radial positions, but not the longitudinal offsets, of the nine doublets are relatively constant with respect to the central pair complex.

To better understand the spatial relationship between the central pair complex and the outer doublets, we performed multibody analysis by treating the central pair protrusions and each doublet as two rigid bodies, refining them separately and remapping the two structures back to each raw subvolume. The spatial relationship of these two rigid bodies in each raw subvolume was then subjected to principal component analysis. For all nine interfaces, we always observed the first principal component, which explains most variations (40-50%), was parallel to the longitudinal axis of axonemes (Supplementary Movie 2), suggesting the doublets and the central pair complex from different tomograms meet at different longitudinal offsets. This is consistent with our consensus average of the entire (9+2) axoneme, where doublet microtubules would be resolved as long as their radial positions are fixed. In the case where longitudinal alignment is heterogeneous, the continuous tube density would be less affected compared to discrete decorating protein complexes including dyneins, radial spokes and N-DRC (Fig. 2d,e).

The multibody analysis also yielded a map with the two rigid bodies placed at their average positions, allowing us to examine the potential ways in which the doublets and central pair complex interact. Interestingly, we observed that most radial spoke heads were separated by a short distance from the central pair protrusions, without any intervening densities (Fig. 3a). However, at the central pair interface with doublet 8, we observed some protrusions from the central pair complex that fit into the “slot” of radial spoke 1 and 2 and also the two openings of the “S-curve” of the radial spoke 3 (Fig. 3b- d). Together, these analyses revealed that radial spokes from the nine doublets are located proximal to nine unique stripes of protrusions from the asymmetric central pair complex with longitudinal variations, with one central pair protrusion showing a classic “train-on-a-track” interaction with the radial spokes head while others resembling the “maglev train system”.

**Figure 3.**
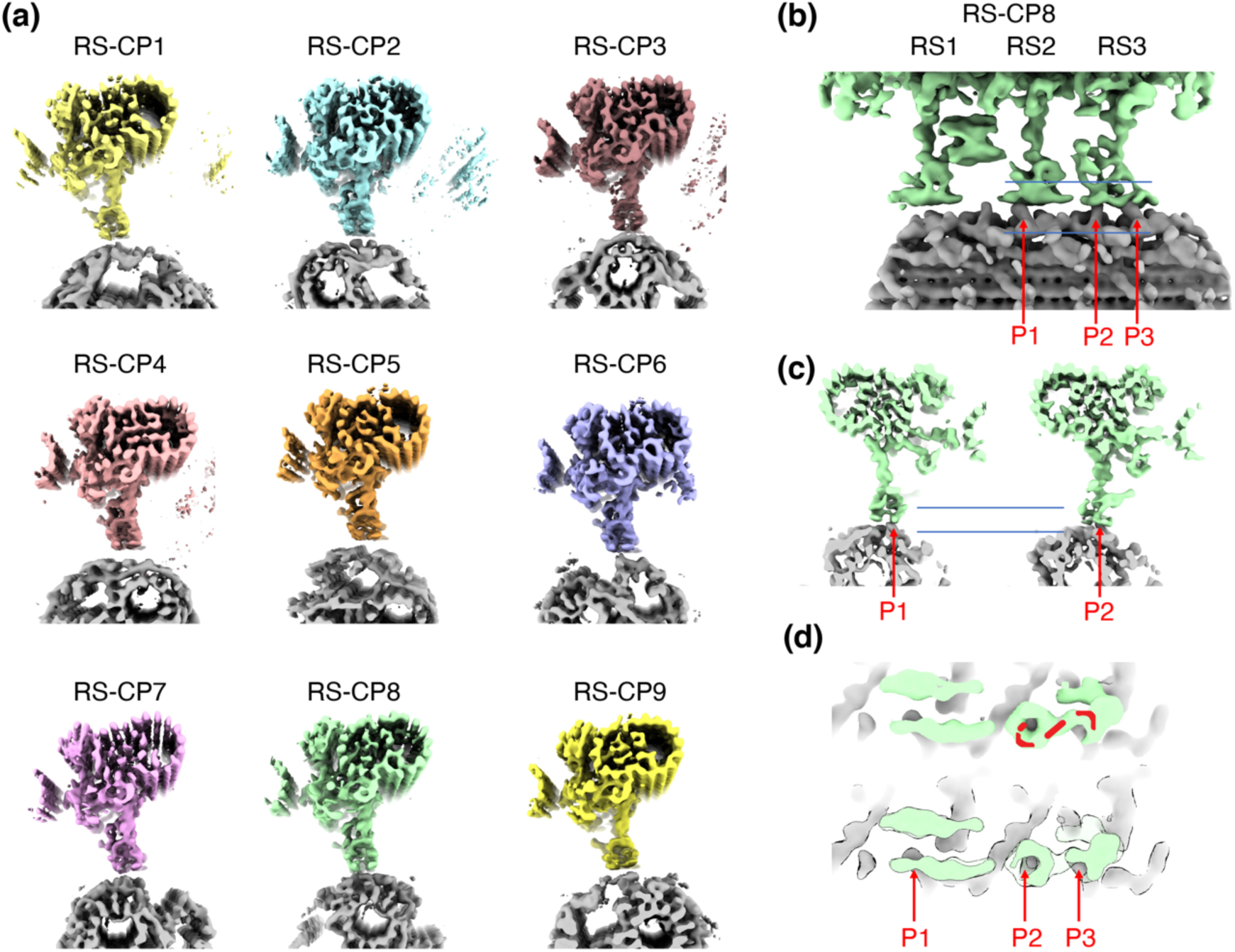
The diverse interfaces between radial spokes and central pair. **a**, The interfaces between the nine radial spokes and the central pair complex are shown and are labeled as “Radial Spoke – Central Pair” and the doublet number (RS-CP1 denotes the interface between radial spokes from doublet 1 and the central pair. Note different proteins from the central pair complex interact with the radial spokes heads from the nine doublets. **b-d**, Three different views of the interface between the radial spokes from doublet 8 and the protrusions, noted as P1, P2, P3, from the central pair complexes. **b**, Longitudinal view of the interface. **c**, Two cross-sections views showing how the RS2 and RS3 from doublet 8 interact with the protrusions of central pair. A section view of the interface is indicated by the two horizontal lines and shown in **d**.

### Asymmetric distribution of sperm-specific features revealed by per-doublet averaging

We then sought to investigate if the outer doublets themselves differ from each other. Based on the contextual information of the nine doublets around the asymmetric central pair complex in our *in situ* cryo-ET data, we grouped subvolumes based on their radial position (numbered as 1 through 9 as in references^28, 29^). The subvolumes were aligned and the averages were calculated for each of the nine positions (N = 800-1000 subvolumes per doublet from 48-55 tomograms). This processing strategy identified unique densities emerging from a location adjacent to the inner arm dyneins from doublet 5 and connecting to multiple positions in doublet 6 (Extended Data Fig. 6a-f). These connecting densities are similar to the “5-6 bridge” observed in sea urchin at lower resolutions^15, 30^, validating our assignment of doublets based on radial positions. Interestingly, the radial spokes and other 96-nm repeating features on both doublet 5 and 6 could be resolved concurrently after local refinement, indicating there is a relatively consistent longitudinal offset (∼20 nm) between these two doublet microtubules throughout different axonemes (Extended Data Fig. 6d). The correlation of the unique bridge densities and the consistent offset that is observed only between doublet 5 and doublet 6 suggest that the bridge could limit the relative sliding between this outer doublet pair (see another example in Extended Data Fig. 6g,h).

Next, we systematically compared motor and non-motor/regulatory protein complexes across the nine doublets. Outer arm dyneins across all nine doublets were indistinguishable from our consensus average (as shown in Figure 1d). The densities of inner arm dyneins were also generally similar for all doublets, with two exceptions. For doublet 5, densities corresponding to dynein c and e (nomenclature defined in *Chlamydomonas*^14, 31^) were shifted compared to the other doublets (Extended Data Fig. 7a), while for doublet 9, densities for dynein b were not resolved (Extended Data Fig. 7b). These results indicate that the motor proteins are largely the same with minor variations in a couple of outer doublets.

We next examined the radial spokes from each doublet. Strikingly, we observed that the sperm-specific features were asymmetrically distributed across the nine doublets (Fig. 4). The barrel density was not observed in doublet 1 and doublet 9 and was present at a lower occupancy in doublet 3. In the remaining six doublets, the occupancy of the barrel was comparable to other repeating structures (e.g. radial spoke 1), suggesting their consistent presence throughout these subvolumes. The RS2-RS3 crosslinkers are absent in doublets 3 and 8, but present in the remaining doublets. In addition, for doublets 2 and 3, we resolved extra “scaffold” densities (RS3 scaffold), close to the base of RS3 (Fig. 4), that were not observed in our consensus map or previously reported consensus maps, in which subvolumes from all nine doublets were averaged together^20, 27^. This is likely because only one or two doublets out of the nine possess these features (11-22%) and averaging did not yield strong signals. Our processing strategy allowed us to separate these subvolumes based on contextual information and the high occupancies of these unique structures in the respective doublets suggest their consistent presence.

**Figure 4.**
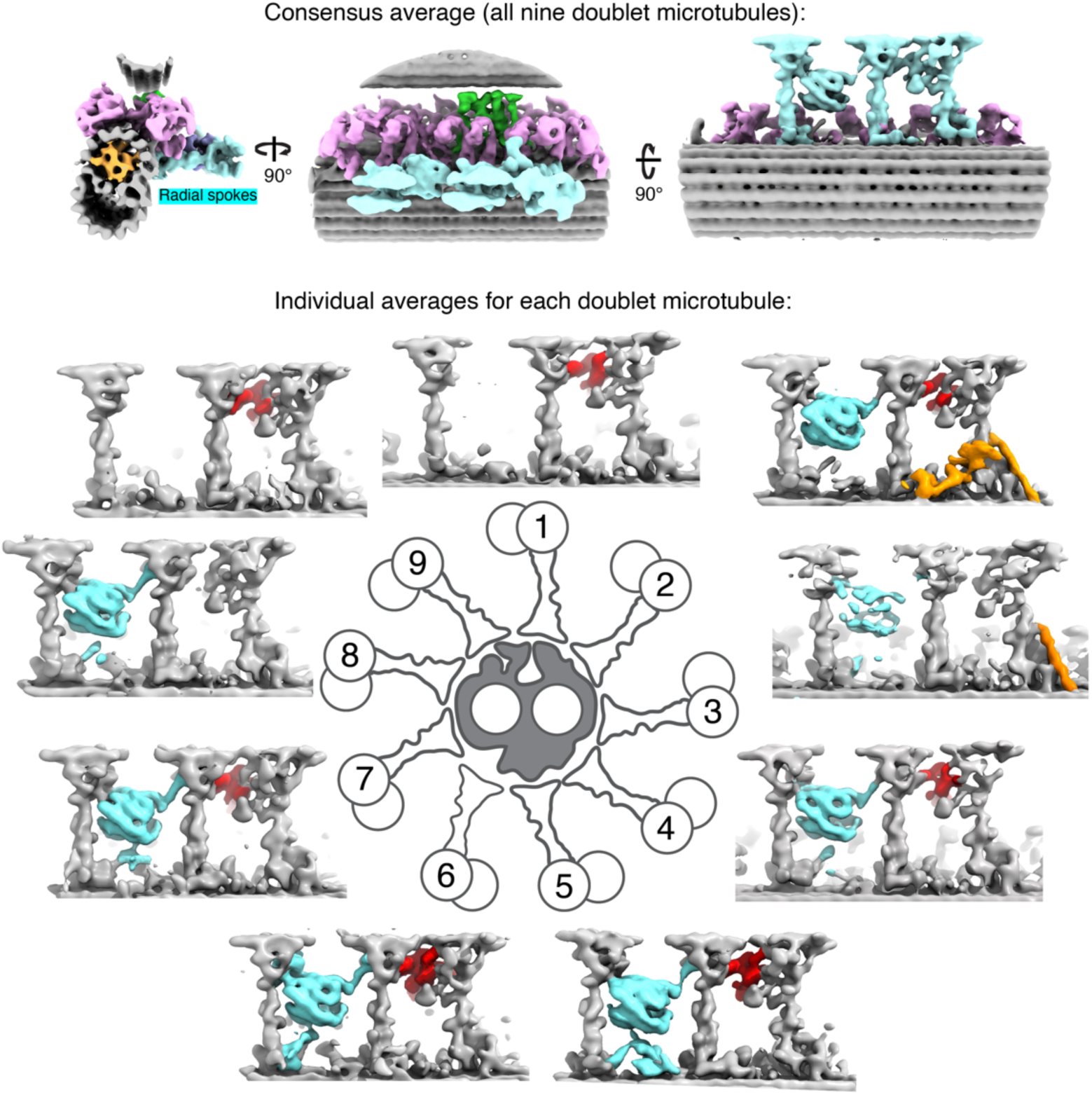
Asymmetric distribution of sperm-specific features in radial spokes from the nine doublet microtubules in mouse sperm. Three orthogonal views of the consensus doublet average of mouse sperm axoneme are shown as in Fig. 1b-d. At the bottom panel, densities corresponding to radial spokes from the nine per-doublet averages are shown around a schematic of a cross-section view of the (9+2)-axoneme. Common features are colored in grey, while the barrel, RS2-RS3 crosslinker and RS3 scaffolds are highlighted in cyan, red and orange, respectively.

We also examined the N-DRCs that crosslink neighboring doublet microtubules (Figure 5). All N-DRCs across the nine doublets share the common “V” shape density, but the extra connections to the microtubule (the “N” structure) show heterogeneities. Interestingly, N-DRCs from doublets 2, 3, 4 share an arch-shaped density perpendicular to the microtubules while the ones from doublets 6, 7, 8 have 45°-tilted thin strands. Doublets 1, 5 and 9 all have extra densities that are distinct from the others, leading to five different sets of N-DRCs. Note all these different features were observed at similar signal-to-noise levels and since they resulted from averaging of more than 800 subvolumes sampled in ∼50 different axoneme tomograms, they represent the commonly shared features within each individual doublet.

**Figure 5.**
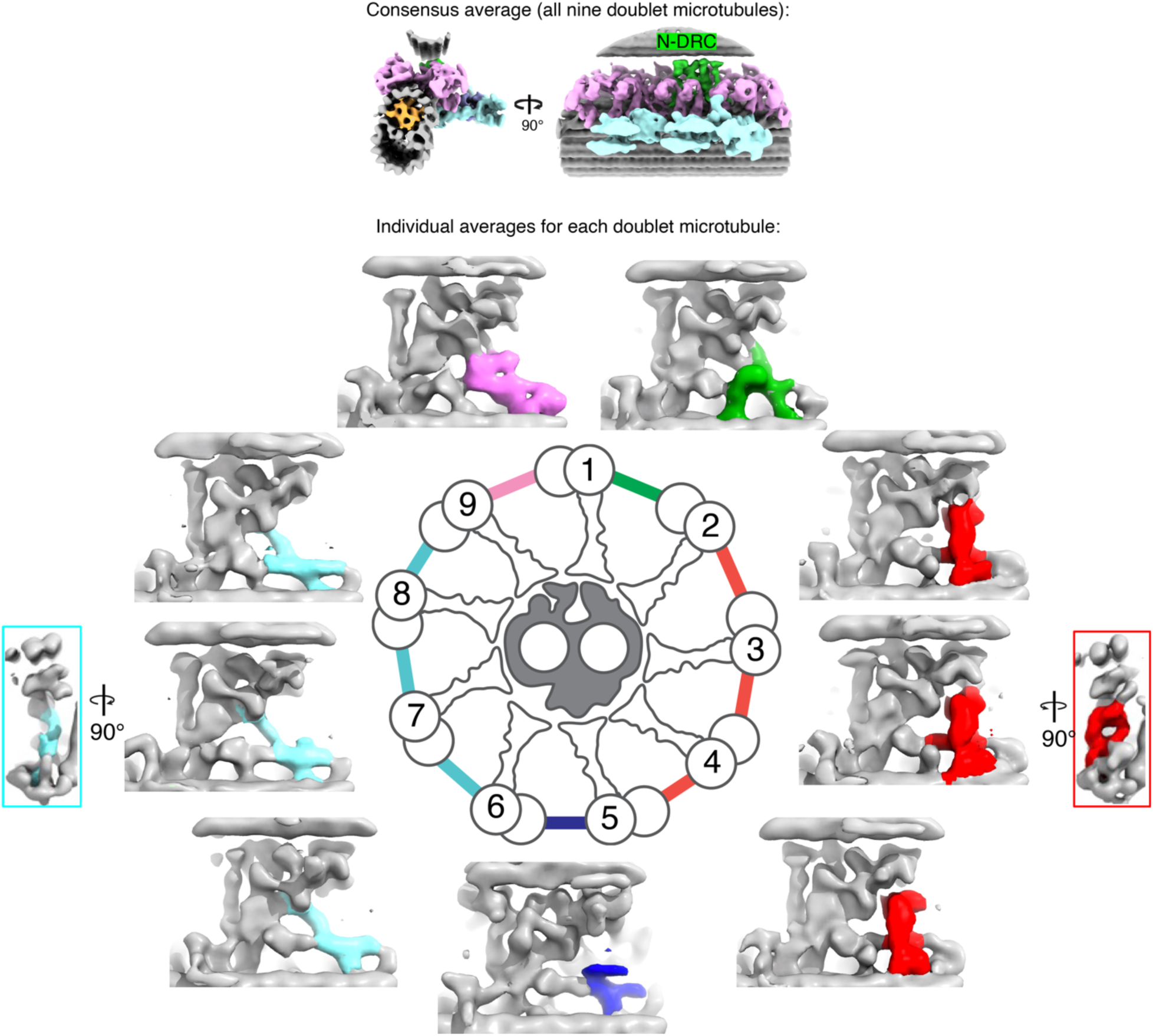
Asymmetric distribution of sperm-specific features in N-DRCs from the nine doublet microtubules in mouse sperm. Two orthogonal views of the consensus doublet average of mouse sperm axoneme are shown at the top panel. At the bottom panel, densities corresponding to N-DRC from the nine per-doublet averages are shown around a schematic of a cross-section view of the (9+2)- axonemes. Common features among the N-DRC are colored in grey, while the unique features are highlighted in various colors.

In order to test if the bending states of the axonemes affect the features of radial spokes and N-DRC, we curated subvolumes from tomograms with and without apparent curvatures and calculated per-doublet averages. The same set of features of radial spokes and N-DRC were observed. In addition, we also collected a dataset of demembraned sperm axonemes (48 tomograms) that are not actively beating. The radial spokes and N-DRCs in these non-motile sperm have the same asymmetric features highlighted in Fig. 4 and Fig. 5, suggesting the asymmetric densities were not caused by bias in macroscopic curvatures, but likely intrinsic features of each doublet.

Our *in situ* data also allowed us to separate axonemes of the midpiece and principal piece based on the presence of mitochondrial and fibrous sheaths, respectively. We averaged subvolumes from these two regions for each doublet and found only subtle differences in the base of RS3 of doublet 2 and also the RS1 of doublet 7 (Extended Data Fig. 8c) (N=400-500, ∼35 Å at FSC=0.143), suggesting the consistency of the axoneme structure along most of the flagella. Overall, our “per-doublet” averaging approach showed that the distributions of various sperm-specific features for both the radial spoke complexes and the N-DRC follow distinct patterns, such that every doublet is decorated by a unique combination of proteins.

### Human sperm share conserved sperm-specific features with different distributions

We next examined whether the unique outer doublet features observed in mouse sperm were also conserved in human sperm. We collected tilt series of intact human sperm (N = 64, not milling lamella), focusing on the thinner principal piece of the flagella and calculated consensus averages of doublet microtubules and the central pair complex (Fig. 6a,b). These consensus maps of human sperm were similar to the ones from mice, with the notable exception that the relative occupancy of the barrel between RS1 and RS2 was much lower in the human sperm axoneme. Using the “per-doublet” processing strategy described above, we then calculated averages for each of the nine doublets individually. Although the resolutions of our averages for the outer doublets from human sperm were lower than those obtained mouse axonemes (∼35 Å vs ∼27Å), they were sufficient to unambiguously identify the aforementioned unique features, including the 5- 6 bridge, the barrel and RS3 scaffolds (all with dimensions >100 Å). In particular, the localization of RS3 scaffolds and the RS2-RS3 crosslinkers show similar asymmetric distributions between human and mouse sperm. However, we observed that the barrel density only existed in four out of nine doublets, in contrast to seven out of nine doublets in mouse sperm axonemes (Fig. 6c). This is consistent with the lower occupancy of barrel densities in our consensus average combining subvolumes from all nine doublets in human sperm. In particular, doublet 2 to doublet 5 appear to be different in human and mouse axoneme in terms of the presence of the barrel, while the rest of the doublets are similar. We also examined the N-DRC from each doublet and found five distinct shapes for the extra density patterned like those in mouse sperm (Extended Data Fig. 9). In summary, our data indicate the asymmetric architecture of axonemes is a general feature of mammalian sperm axonemes, although there are intriguing variations of distribution for the barrel between mouse and human.

**Figure 6.**
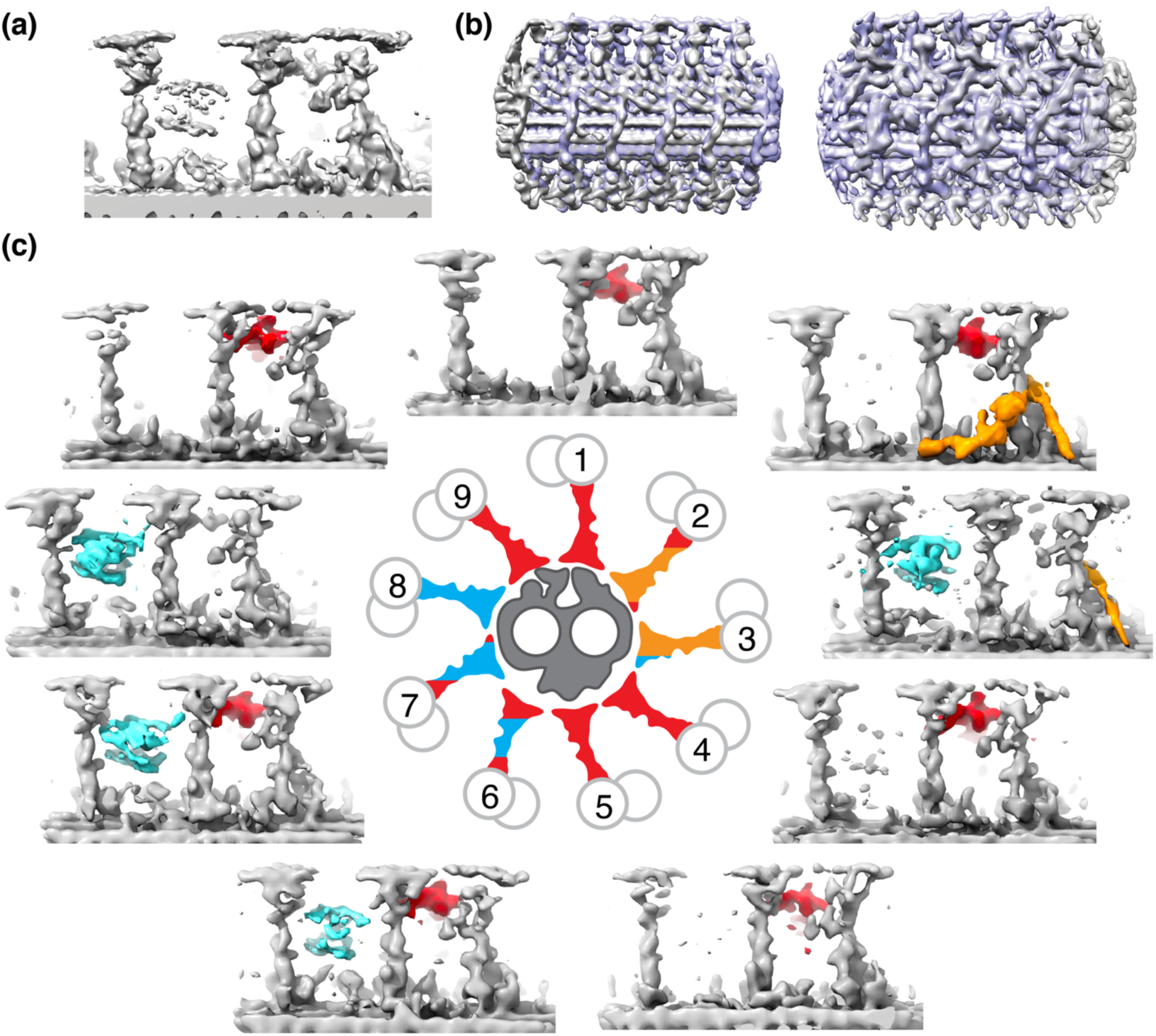
Human sperm have the barrel structures differently distributed in the nine doublet microtubules compared to mouse sperm. **a**, A consensus average map of the radial spoke region for human sperm axonemes. The barrel density appears to have lower occupancy compared to the radial spokes, which is not the case in the consensus average of mouse sperm doublets in Fig. 1g. **b**, The consensus average map of the central pair complex for human sperm axonemes (grey) is overlaid with the one from mouse sperm axonemes (blue). **c**, Densities corresponding to radial spokes from the per-doublet averages are shown around a schematic of a cross-section view of the (9+2)-axonemes. Common features among the radial spokes are colored in grey, while the barrel, RS2- RS3 crosslinker and RS3-base features are highlighted in cyan, red and orange, respectively.

## Discussion

Our *in situ* tomography studies of mouse and human sperm revealed a large ensemble of macromolecular complexes in their native cellular environment. In particular, we observed various sperm-specific features in the MIPs, radial spokes and N-DRC that were not observed in mammalian respiratory cilia or non-mammalian sperm. Furthermore, we reconstructed the entire axoneme using contextual information and uncovered the asymmetric architecture of mammalian sperm axoneme, where every microtubule-based structure is decorated by a unique set of non-motor proteins. As these non-motor proteins regulate the dynein activity based on previous studies, we propose that the asymmetries of non-motor protein complexes could modulate the sliding of the nine doublets individually to shape the species-specific asymmetric waveforms observed for mammalian sperm, as discussed below (Fig. 7a,b).

**Figure 7.**
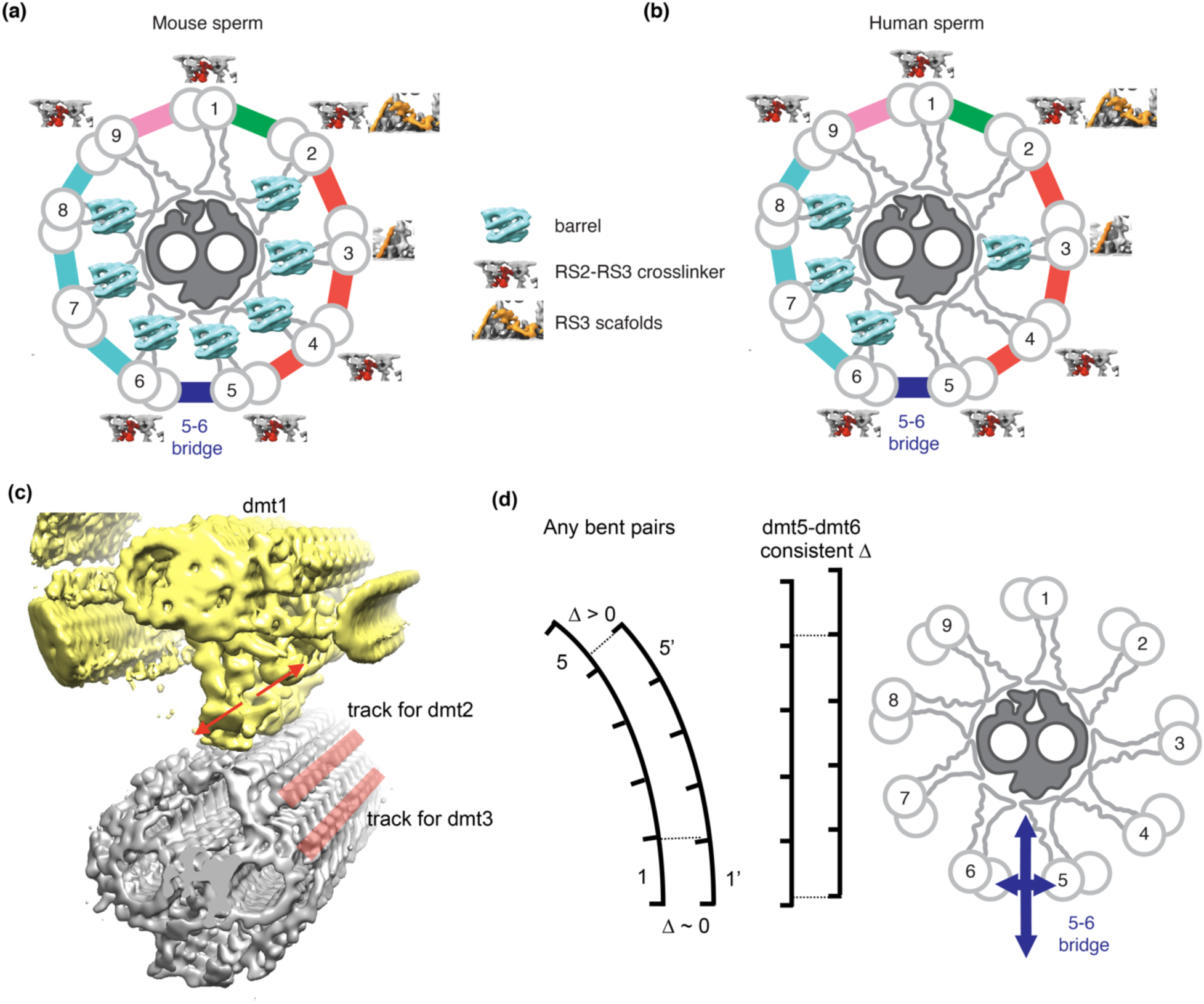
Model: every outer doublet is surrounded by a different set of regulatory complexes in mammalian sperm. Schematic of the (9+2) axonemes of mouse (**a**) and human sperm (**b**). The doublets are numbered and the sperm-specific regulatory complexes are labeled for each of the nine outer doublets. In particular, the components from the radial spokes are shown for each doublet and the N-DRCs are colored differently depending on the extra density (as in Fig. 5). Note the barrel distribution is different in mouse and human sperm axonemes. **c**, a model consistent with the data: radial spokes from different doublets interact with specific stripes of protrusions from the central pair. **d**, Schematics showing gradual accumulations of offsets along bending curves (left panel) and likely situation for doublet 5-doublet 6 (middle panel). The consistent offset would be consistent with limited horizontal bending as shown (right panel).

### Sperm-specific structures added additional mechanical coupling

Mammalian sperm flagellar are generally much longer and wider compared to flagella from unicellular organisms or respiratory cilia (lengths of >45 µm *vs.* ∼10 µm and diameters of > 0.5-1 µm *vs.* ∼0.3 µm). Sperm axonemes are also surrounded by additional subcellular structures, such as outer dense fibers, mitochondrial and fibrous sheaths, which likely present additional mechanical challenges to beat generation. Previous studies also suggested larger bending torques are associated with mammalian sperm compared to other cilia^32^, but it has remained unclear where sperm axonemes have evolved specific mechanisms to withstand the additional mechanical stress. Our averages of *in situ* tomography data revealed many additional non-motor proteins that crosslink the known axonemal components, including the MIPs that connect all thirteen protofilaments of the A-tubule of the doublets, the barrel and RS2-RS3 crosslinker between the three radial spokes, the scaffolds at the base of radial spokes, and extra densities in addition to the common “V”-shaped N-DRC. We propose that these additional proteins function to strengthen the mechanical rigidity of the corresponding components to accommodate higher mechanical requirements of sperm axonemes. As the microtubules must be able to withstand the bending and the seam is the weakest point^33, 34^, the extensive MIP network that we observed, with proteins stapling across the A9 and A10 protofilaments, might be expected to lead to greater stability. The radial spokes were previously observed to tilt relative to the microtubule^30^. Additional connections between the three radial spokes and the scaffolds at the bases would integrate them together into a more rigid unit. Another possibility is that the coupling could lead to coordinated movement of the three radial spokes. The N-DRCs regulate the sliding between the neighboring doublets and prevent splaying of axonemes^10^. Extra densities linking to the microtubules could improve the stability of N-DRC under higher mechanical stress. Together, the additional protein complexes in mammalian sperm would help to maintain the geometric integrity of the (9+2)-microtubule configuration of the axoneme under mechanical stress during beating.

### The spatial relationship between the nine doublets and the central pair complex

Previous studies suggested that the central pair can twist radially relative to the nine outer doublets in *Chlamydomonas* flagella^35^. Our observation of densities corresponding to the nine doublet microtubules in the average of the entire (9+2) axoneme (Fig. 2d,e) can exclude the possibility of free twisting of the central pair to the doublets in mammalian sperm as the doublet densities would be smeared by averaging. Interestingly, we observed a cleft between the two halves of radial spokes 1 and 2 and holes in the radial spoke 3 in mammalian sperm axonemes (Fig. 1c), in contrast to the flatter surface of the radial spoke heads from *Chlamydomonas* flagella^11–13^. While the flat surfaces may enable the twisting in *Chlamydomonas*^35^, the complementary shapes of radial spokes and protrusions from the central pair complex in mammalian sperm may restrict such radial movements (as shown in Fig. 3). Fixation of radial positions of the nine doublets also positions each of the radial spoke complexes in proximity to unique stripes of densities of the central pair complex (Fig. 7c). Together, this arrangement could allow functional specialization and divergent evolution of each doublet microtubule, such as the distinct sets of sperm-specific features in the nine doublets.

When subvolumes of the central pair were expanded and averaged (Fig. 2d,e), the low occupancies of densities corresponding to the periodic and discrete features, including the radial spokes, dyneins and N-DRCs, suggest the lack of alignment along the longitudinal direction. This observation could be explained by heterogeneity in the longitudinal offsets between doublets and the central pair complex in the randomly sampled axoneme positions (N=66) that were combined in the averages. This idea is consistent with the sliding hypothesis for axonemal bending, in which active dyneins generate displacement between the neighboring doublets and lead to relative movement relative to the central pair complex at the same time. Consistent with this idea, most radial spoke/central pair interfaces do not possess structurally consistent densities in between, allowing free longitudinal sliding of nine doublets on the nine tracks on the central pair complex (Fig. 3).

The 5-6 bridge is a sperm-specific feature that appears to be conserved between sea urchin^15^ and mammalian sperm. Our data revealed that the register of doublet 5 and 6 has a consistent offset (Extended Data Fig. 6), based on averages of subvolumes sampled from random positions or cells of mouse sperm. Previous studies pointed out for two parallel inelastic microtubule filaments, bending would lead to gradual accumulation of longitudinal offsets between if they are at different radii of the bend since the outer curve will be longer than the inner one (Fig. 7d) ^30^. The consistent offset between doublet 5 and 6 suggests that there is limited bending along the direction parallel to the plane of the two filaments. The bundling of these two doublets can create a stiffer filament compared to all other doublets, with distinct elastic properties or bending propensities in different directions. Such asymmetries of mechanical properties within the nine doublets could also contribute to complicated waveforms.

### Asymmetric distribution of regulatory complexes in sperm axonemes may lead to asymmetric beating waveforms

Our *in situ* data and processing strategies based on the contextual information revealed that the axoneme itself, which appeared to have “pseudo nine-fold symmetry” in the classical TEM images^3^, is highly asymmetric at the molecular level. As the axoneme is the underlying engine that drives the flagellar beating motion, such asymmetries in structure could lead to asymmetric beating waveforms.

Furthermore, the asymmetries lie mostly in the mammalian sperm-specific non-motor protein complexes, including the radial spokes and N-DRCs. These complexes are well-established regulators of dynein motor activities and defects in individual protein components can lead to irregular beating^7, 8, 10, 11^. In mammalian sperm, we show here that each of the nine doublet microtubules is decorated by a unique composition of these regulators. We speculate that this distinct molecular composition could lead to differences in the sliding speeds, or bending forces for each of the doublet pairs (Fig. 7). The non-equivalent forces could lead to a deviation from the single plane of beating characteristic of more symmetric axonemes. Thus, we hypothesize that the non-uniform distribution of sperm-specific radial spoke and N-DRC regulators are important for asymmetric and non-planar beating. Additionally, previous studies reported that human and murine sperm have different swimming waveforms^19^. Our studies suggest that although the radial spoke barrels are conserved in human and mouse, their distribution varies and this variation could create diversity in sperm swimming behavior.

This study has revealed mammalian sperm-specific structures within the radial spokes and N-DRCs. The molecular identity of these structures remains unknown and constitutes an important problem for future investigation. The still unknown proteins likely arose to serve functions required by natural fertilization in mammals. Sperm from different species are cast into a foreign environment and selected for their ability to reach and fertilize an egg. Sea urchin and zebrafish sperm swim in water whereas mammalian sperm swim in a thin layer of viscous liquid on uneven surfaces of the female reproductive tracts^36^. Asymmetric and non-planar waveforms of mammalian sperm could be beneficial to navigate through viscous environments and around three-dimensional obstacles. Furthermore, the dimensions and physical characteristics of reproductive tracts vary among different mammals, so fine-tuning of the waveforms, and the underlying structural features to produce distinct waveforms, are likely under strong evolutionary selection. In the future, systematic genetic analyses of mammalian sperm-specific proteins would be valuable to connect sperm-specific axonemal structures with their functions in sperm motility and reproductions.

## Acknowledgement

We thank Shixin Yang from the CryoEM facility at Janelia Research Campus for their assistance with data collection. We thank Zanlin Yu, Eric Tse and David Bulkley in the UCSF EM core facility for their assistance with data collection. We thank Sam Li and Shawn Zheng at UCSF for suggestions on EM data processing. We thank Tom Goddard at UCSF for providing a script to mark coordinates of subtomograms. EM data processing utilized computing resources at both the workstations at Janelia Research Campus and HPC Facility at UCSF. We are grateful to members of the Vale laboratory and Agard laboratory for discussions and critical reading of the manuscript. Z.C. was supported by the Helen Hay Whitney Foundation Postdoctoral Fellowship. P.V.L. received funding from Pew Biomedical Scholars Award and GCRLE grant from Global Consortium for Reproductive Longevity and Equality made possible by the Bia-Echo Foundation. D.A.A. received funding from NIH R35GM118099. R.D.V. received funding from NIH R35GM118106 and the Howard Hughes Medical Institute.

## Author contributions

Z.C., G.A.G. and R.D.V. conceived the project and designed the experiments after discussions with other authors. S.Z. and C.G. provided the mouse sperm sample. M.S. and Z.C. prepared mouse sperm lamellae grids. W.S. and P.V.L. provided human sperm samples. Z.C. and Y.L. prepared the human sperm grids. M.S., Z.C. and G.A.G. performed the FIB-SEM processing. Z.X. and S. Y. optimized the data collection. Z.C. processed the data with help from M.S., X.Z and R.Y. and suggestions from G.A.G. and D.A.A.. Z.C. and R.D.V. wrote the manuscript draft with comments from all authors.

## Data Availability

The cryoEM maps of the mouse sperm axonemes have been deposited in the Electron Microscopy Data Bank under the accession codes EMD-XXXX (will be submitted before publication). All data analyses are included in this Article and the Supplementary Information. Other relevant data are available from the corresponding author upon requests.

---

**Extended Data Fig. 1.**
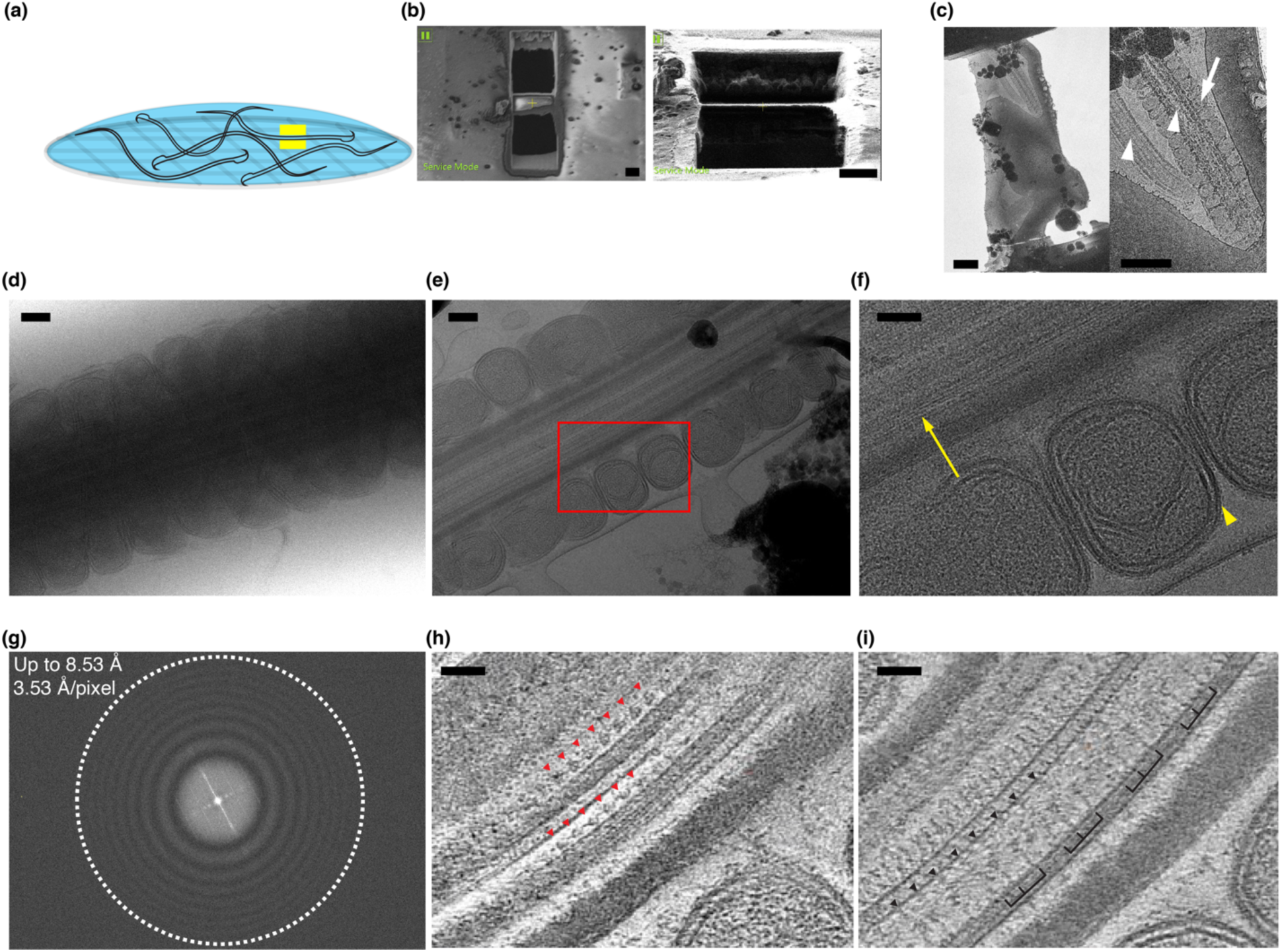
The workflow of FIB-SEM/cryo-ET and representative images. **a**, Schematic of EM grids with vitrified mouse sperm. An example for where the windows were milled by FIB-SEM is denoted by two yellow boxes. **b**, SEM (left panel) and FIB (right panel) images of a lamella are shown (scale bar: 5 µm). **c**, A low-magnification montage and its zoom-in view of a lamella acquired in 300 kV Titan Krios microscope (scale bar: 1 µm). Axoneme and mitochondria are indicated by white arrowheads and an arrow, respectively. Note there are two flagella, one with the mitochondrial sheath and the other with the fibrous sheath, corresponding to the midpiece and the principal piece of the flagella, respectively. **d**, A representative image of non-milled sperm axoneme recorded by 300 kV Titan Krios microscope at a total dose of 3 e^-^/Å^2^ (scale bar: 100 nm). **e**, A representative image of milled lamella (thickness ∼300 nm) is shown (scale bar: 50 nm). The zoom-in view in **f** is outlined using the red rectangle. **f**, Individual protofilaments of microtubules and bi-layered membranes of the mitochondria are indicated by yellow arrows and arrowheads. Fourier transform of the image shown in **e** with the upper limit of resolution when the thong rings are fitted using Ctffind4. Note the limit is 8.53 Å, which is close to the Nyquist limit at the pixel size of 3.53 Å. **h**, **i**, Two sections of 3D tomograms are shown. Repeating outer arm dyneins (every 24 nm) anchored on doublet microtubules are indicated using red arrowheads in **h**. Repeating protrusions (every 32 nm) extending from the singlet microtubule are indicated using black arrowheads in **i**. Repeating patterns of three radial spokes on the doublet microtubules (every 96 nm) are grouped and indicated using black brackets in **i** (scale bar: 50 nm).

**Extended Data Fig. 2.**
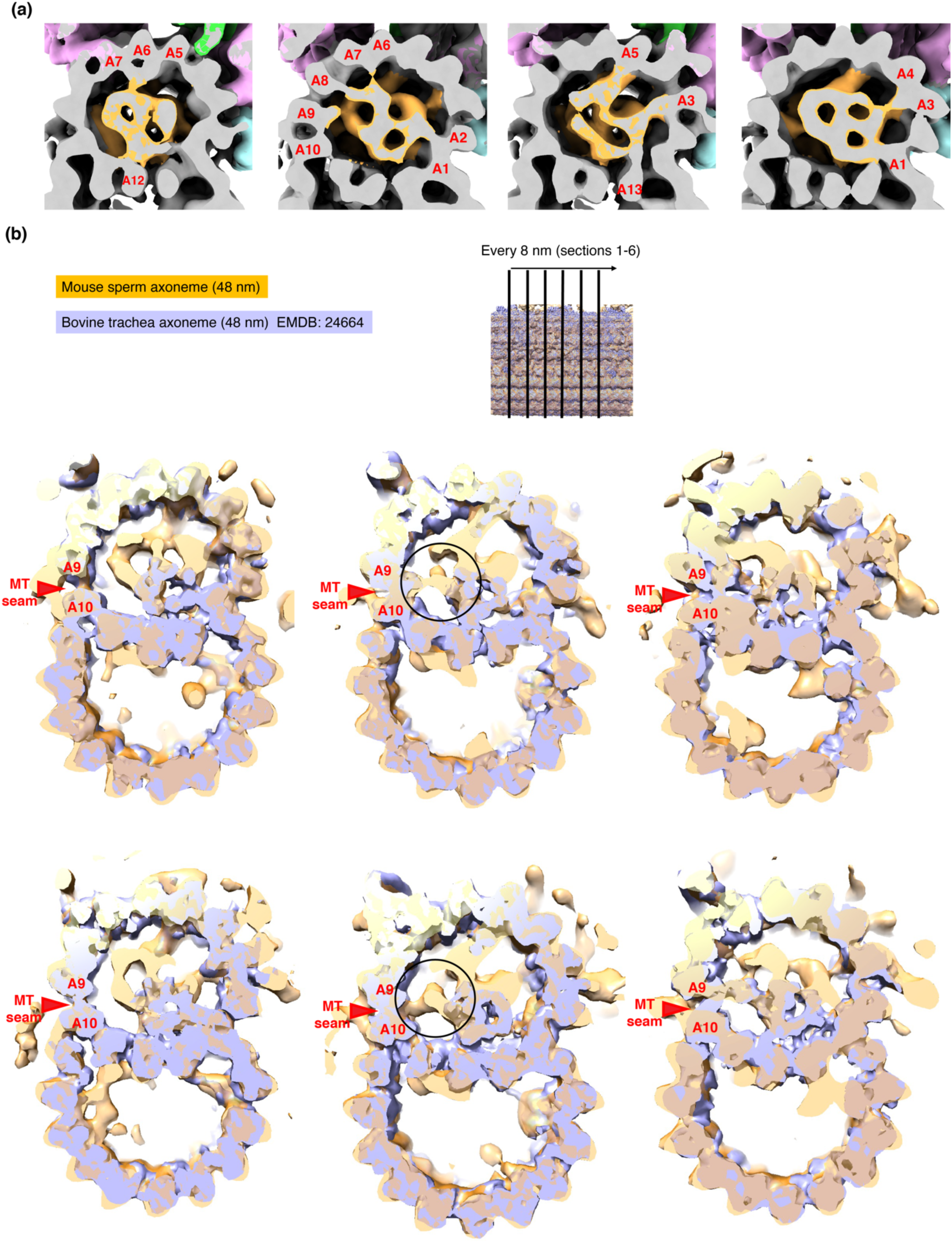
MIPs network in A-tubule of doublet microtubules. **a**, Four cross-section views of the A tubule in the doublet microtubule are shown. The MIPs network is colored in orange. Three modes of interactions between the MIPs and the protofilaments are observed (see text). The protofilaments connecting to the MIPs are also labeled. **b**, Six cross-section views of the overlay of the average of the 48-nm repeat of doublets of mouse sperm (orange, this study) and the 48-nm repeat of doublets of bovine trachea cilia (blue, EMDB: 24664). The contour levels of the two maps are adjusted so that the densities of microtubules are matched. The microtubule seam, A9 and A10 protofilaments are labeled in all views. Note there are additional densities near A9-A10 region in the mouse sperm axoneme.

**Extended Data Fig. 3.**
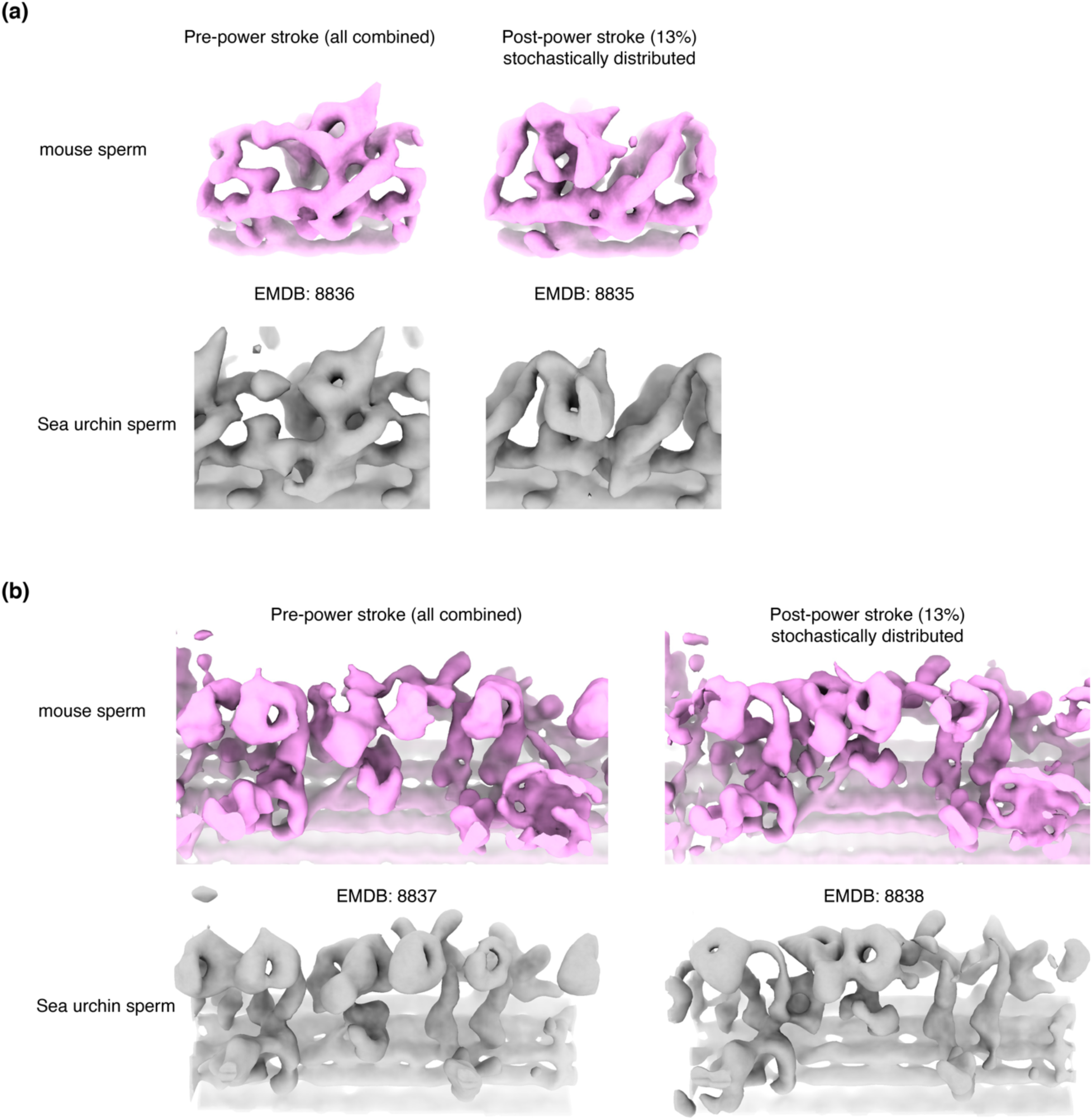
Axonemal dyneins in mouse sperm axonemes. **a**, Comparison of outer arm dyneins in mouse sperm axoneme (this study) and published maps for those from sea urchin sperm (EMDB: 8835 and EMD: 8836). **b**, Comparison of inner arm dyneins in mouse sperm axoneme (this study) and published maps for those from sea urchin sperm (EMDB: 8837 and EMD: 8838).

**Extended Data Fig. 4.**
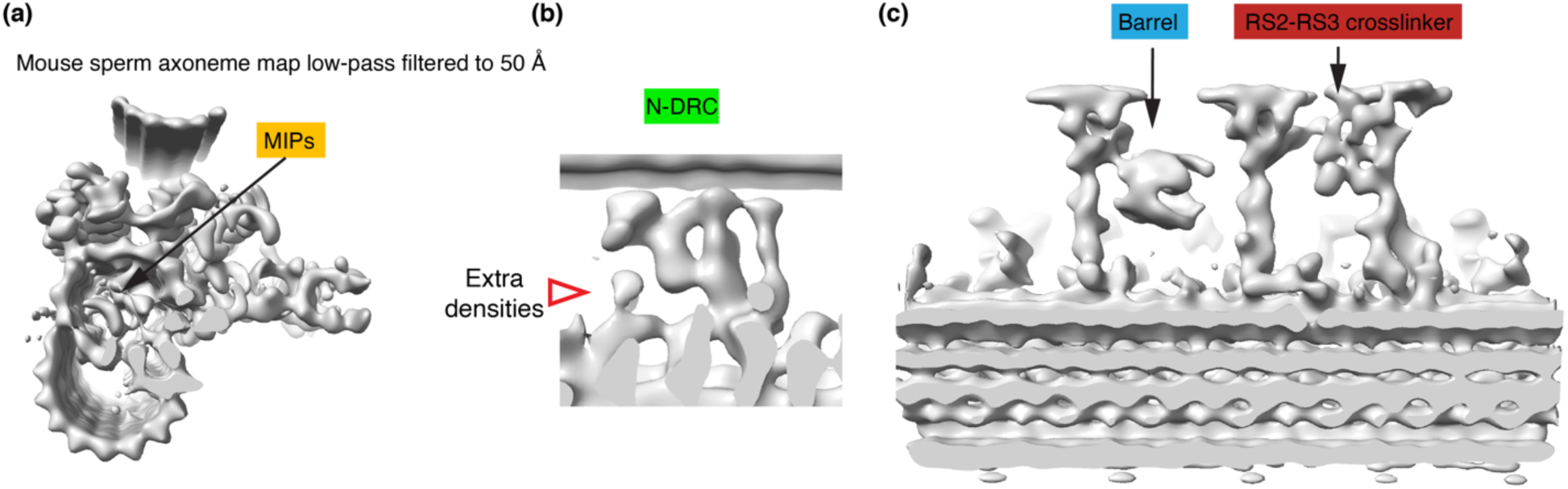
Sperm-specific features are apparent in a low-pass filtered map (to 50 Å) for the mouse sperm axonemes. **a-c**, The doublet average of mouse sperm axoneme is low-pass filtered to 50 Å and viewed from different angles. Note this resolution is lower than published axoneme maps (e.g. 34Å for EMDB: 5950 shown in Fig. 1 for comparison). The extensive network of MIPs (**a**), extra densities at N-DRC (**b**) and radial spokes (**c**) remain apparent while comparing to human respiratory cilia, suggesting the presence of novel densities in our maps are not due to higher resolution, but most likely the presence of additional proteins.

**Extended Data Fig. 5.**
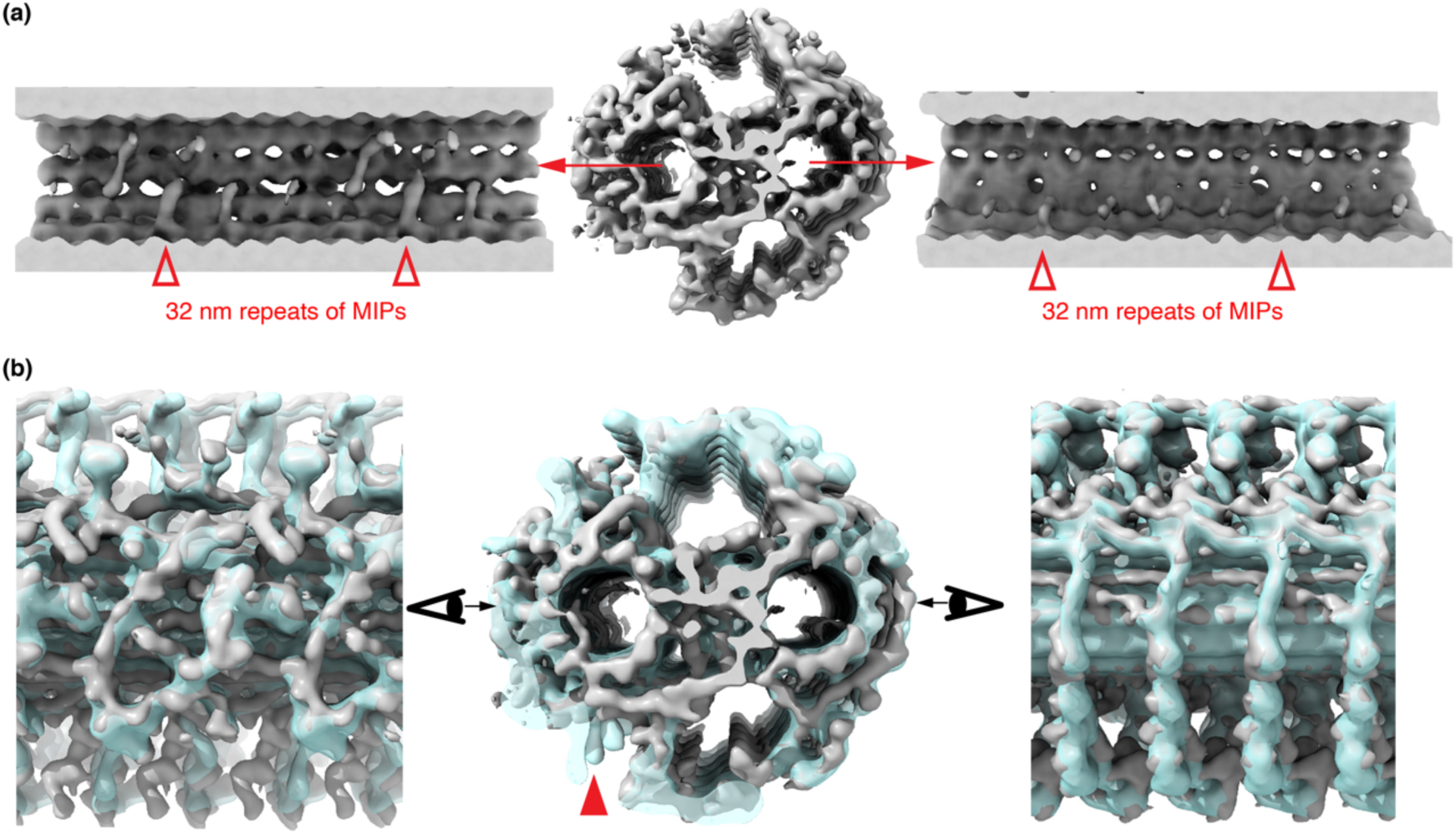
The central pair complex in mouse axoneme possesses more MIPs but the overall shape is similar to the one from sea urchin sperm. **a**, Two longitudinal views of the MIPs inside the two singlet microtubules are shown. Periodic features (every 32 nm) are indicated by the red empty arrowhead. **b**, The average map of the central pair complex in mouse sperm axonemes is overlaid on the published map for the central pair complex from sea urchin sperm (EMDB: 9385) (REF) and they are colored in grey and cyan, respectively. Note most of the protrusions are very similar and one difference is indicated by the solid red arrowhead in the middle panel.

**Extended Data Fig. 6.**
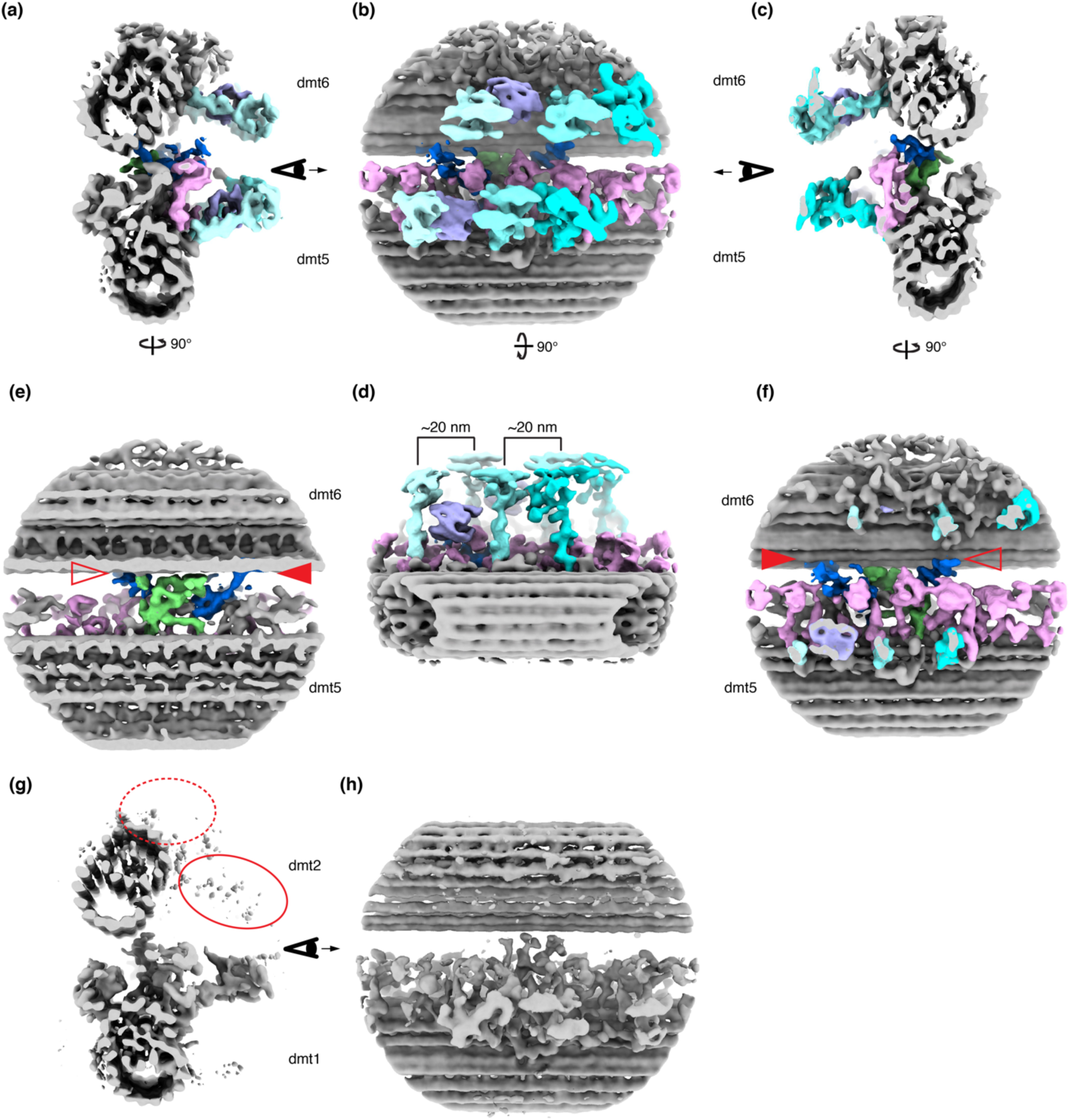
The 5-6 bridge of mouse sperm axonemes. **a-d**, Four views of the composite map for 5-6 bridge in mouse sperm axonemes combining maps for local refinement of the doublet 5, doublet 6 and the bridge. The inner arm dyneins, radial spokes, the barrel, N-DRC and extra bridge densities are colored in pink, different shades of cyan, light blue, green and blue, respectively. Note that the radial spokes which repeat every 96 nm on both doublet 5 and doublet 6 are resolved and there appears to be a ∼20 nm offset between their longitudinal registers, as indicated in the staggering distance of RS1 and RS2 from doublet 5 and doublet 6 in **d**. **e**, **f**, Two cut-in views of **a** and **c** are shown. Bridge-specific densities are highlighted in blue. These densities extend from the inner arm dyneins towards doublet 6. Direct contacts of the bridge-specific densities to doublet 6 is indicated by the solid red arrowhead while densities in close proximity of the base of radial spoke 2 in doublet 6 are indicated by the empty red arrowhead. Note the N-DRC from doublet 5 appears connected to doublet 6 (also see Extended Fig. 5). **g**, **h**, Subvolumes for doublet 1 was aligned similarly and equivalent views of **a** and **b** are shown. The positions for dyneins and radial spokes on doublet 2 are circled by dashed and solid red ovals and note they are not resolved even after local refinement.

**Extended Data Fig. 7.**
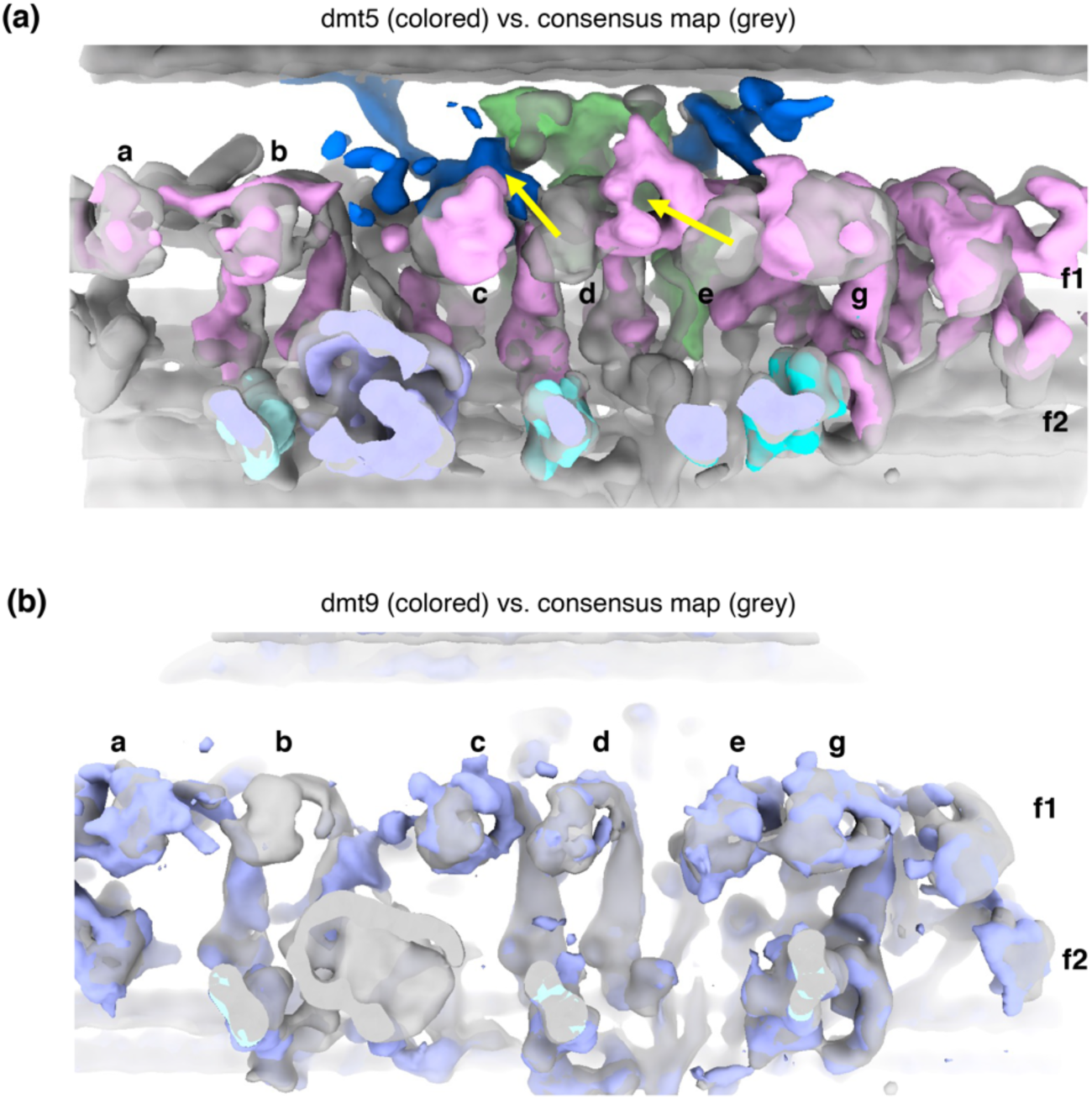
The inner arm dyneins in doublet 5 and doublet 9. **a**, An overlay of inner arm dyneins from doublet 5. The assigned inner arm dyneins, N-DRC and 5-6bridge densities from the doublet 5 are colored in pink, green and blue, respectively while the densities for the consensus map are in grey. Different inner arm dyneins are named according to the nomenclature defined by studies of *Chlamydomonas* flagella. The shifted positions of dynein d and dynein e are indicated by the two yellow arrows. **b**, An overlay of inner arm dyneins from doublet 9 and the consensus average were blue and grey, respectively. Note dynein b in doublet 9 is not resolved.

**Extended Data Fig. 8.**
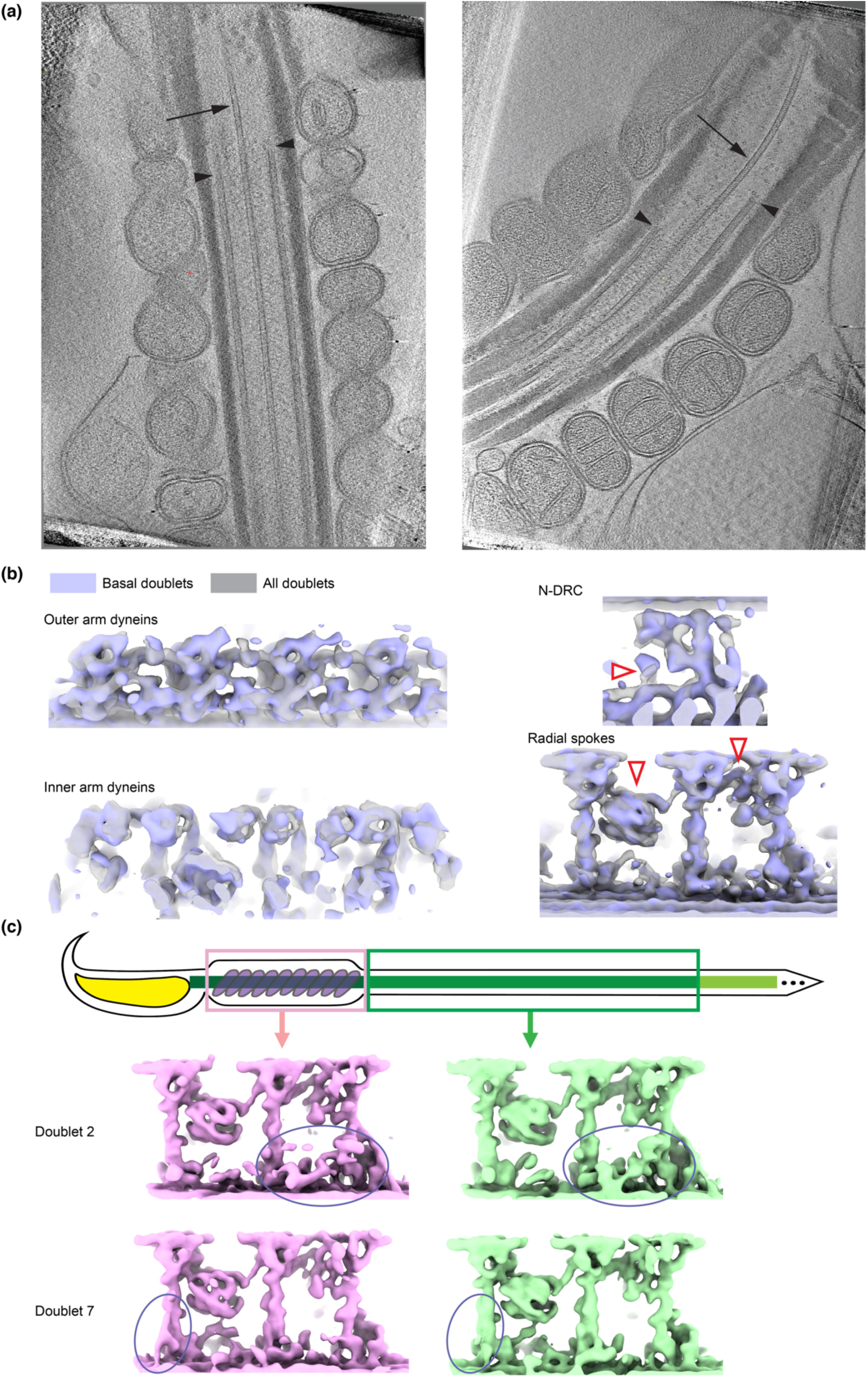
Longitudinal consistency of doublet structures. **a**, Two z-sections of 3D tomograms of the basal region of mouse sperm axoneme are shown. The singlet microtubules extending further into the cell bodies and the beginning of the doublet microtubules are indicated by the arrows and arrowheads, respectively. **b**, Densities corresponding to different protein complexes from subvolume averages of the basal region and all doublet microtubules are overlaid and colored in blue and grey, respectively. Note the sperm-specific features in N-DRC and radial spokes exist in the average of doublet microtubules in the basal region and are highlighted using empty red arrowheads. **c**, Schematic of a mouse sperm flagella with the midpiece and principal piece highlighted in the top panel. Subvolume averages of doublet 2 and doublet 7 from the midpiece and the principal piece are shown. The shapes of densities for the scaffolds at the base of radial spoke 3 appear different in doublet 2 and are circled. Additional densities at the base of radial spoke 1 in the doublet 7 in the midpiece region are observed and circled.

**Extended Data Fig. 9.**
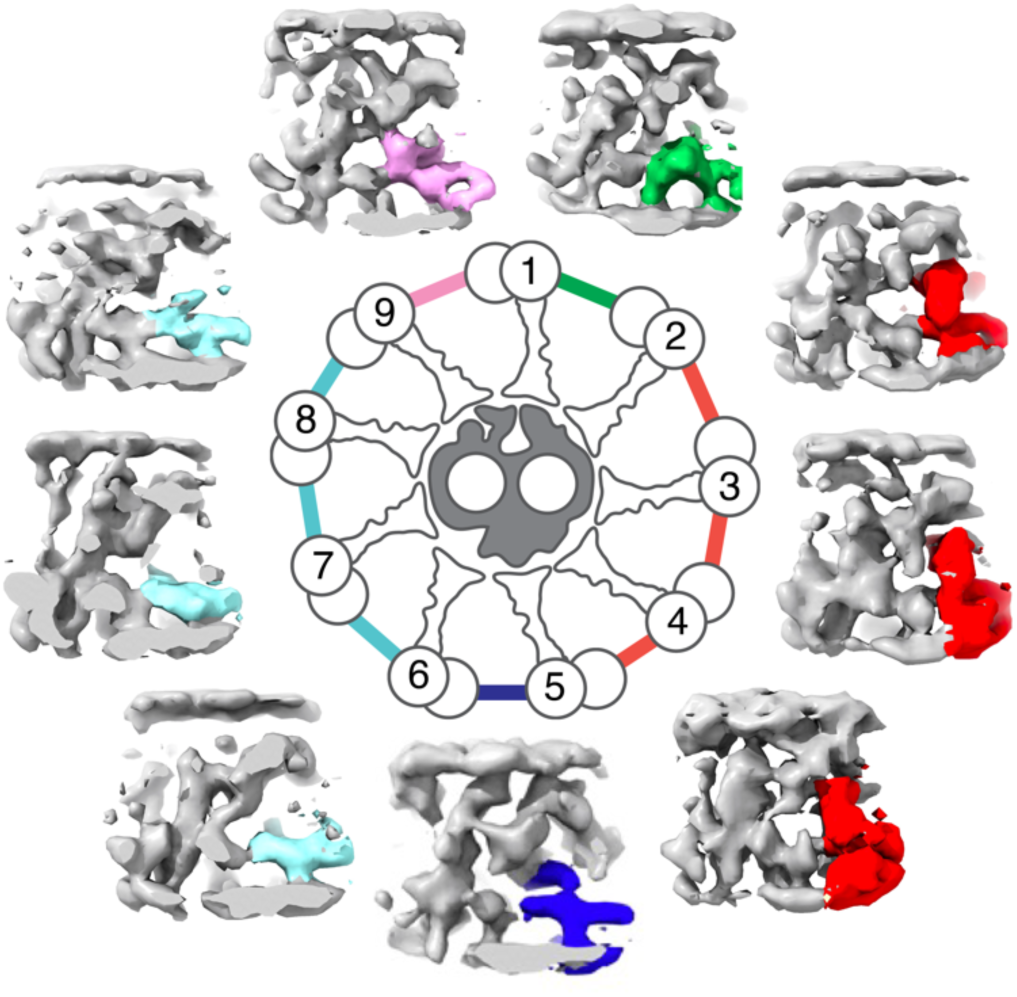
Asymmetric distribution of sperm-specific features in N-DRCs from the nine doublet microtubules in human sperm. Densities corresponding to N-DRC from the nine per-doublet averages are shown around a schematic of a cross-section view of the (9+2)- axonemes. Common features among the N-DRC are colored in grey, while the unique features are highlighted in various colors.

**Supplementary Movie 1. The structure of the radial spoke barrel.**

**Supplementary Movie 2. Multibody analysis of radial spokes from doublet 6 and central pair surface.** The movie contains 10 frames and each frame represents an average of 1/10 of the total populations distributed along the first component of the Principal Component Analysis. The different frames indicate that the radial spokes and the central pair complex can come together at different longitudinal offsets. The interface between doublet 6 and the central pair complex was used as an example.

## References

1. Wallmeier, J., et al. Motile ciliopathies. Nat Rev Dis Primers 6, 77 (2020).

2. Bayless, B.A., Navarro, F.M. & Winey, M. Motile Cilia: Innovation and Insight From Ciliate Model Organisms. Front Cell Dev Biol 7, 265 (2019).

3. Fawcett, D.W. The mammalian spermatozoon. Dev Biol 44, 394–436 (1975).

4. Ishikawa, T. Axoneme Structure from Motile Cilia. Cold Spring Harb Perspect Biol 9(2017).

5. Satir, P., Heuser, T. & Sale, W.S. A Structural Basis for How Motile Cilia Beat. Bioscience 64, 1073–1083 (2014).

6. Lindemann, C.B. & Lesich, K.A. The many modes of flagellar and ciliary beating: Insights from a physical analysis. Cytoskeleton (Hoboken) 78, 36–51 (2021).

7. Witman, G.B., Plummer, J. & Sander, G. Chlamydomonas flagellar mutants lacking radial spokes and central tubules. Structure, composition, and function of specific axonemal components. J Cell Biol 76, 729–47 (1978).

8. Huang, B., Piperno, G., Ramanis, Z. & Luck, D.J. Radial spokes of Chlamydomonas flagella: genetic analysis of assembly and function. J Cell Biol 88, 80–8 (1981).

9. Smith, E.F. & Sale, W.S. Regulation of dynein-driven microtubule sliding by the radial spokes in flagella. Science 257, 1557–9 (1992).

10. Bower, R. et al. The N-DRC forms a conserved biochemical complex that maintains outer doublet alignment and limits microtubule sliding in motile axonemes. Mol Biol Cell 24, 1134–52 (2013).

11. Grossman-Haham, I. et al. Structure of the radial spoke head and insights into its role in mechanoregulation of ciliary beating. Nat Struct Mol Biol 28, 20–28 (2021).

12. Gui, M. et al. Structures of radial spokes and associated complexes important for ciliary motility. Nat Struct Mol Biol 28, 29–37 (2021).

13. Pigino, G. et al. Cryoelectron tomography of radial spokes in cilia and flagella. J Cell Biol 195, 673–87 (2011).

14. Bui, K.H., Yagi, T., Yamamoto, R., Kamiya, R. & Ishikawa, T. Polarity and asymmetry in the arrangement of dynein and related structures in the Chlamydomonas axoneme. J Cell Biol 198, 913–25 (2012).

15. Lin, J., Heuser, T., Song, K., Fu, X. & Nicastro, D. One of the nine doublet microtubules of eukaryotic flagella exhibits unique and partially conserved structures. PLoS One 7, e46494 (2012).

16. Dutcher, S.K. Asymmetries in the cilia of Chlamydomonas. Philos Trans R Soc Lond B Biol Sci 375, 20190153 (2020).

17. Muschol, M., Wenders, C. & Wennemuth, G. Four-dimensional analysis by high-speed holographic imaging reveals a chiral memory of sperm flagella. PLoS One 13, e0199678 (2018).

18. Hansen, J.N., Rassmann, S., Jikeli, J.F. & Wachten, D. SpermQ(-)A Simple Analysis Software to Comprehensively Study Flagellar Beating and Sperm Steering. Cells 8(2018).

19. Babcock, D.F., Wandernoth, P.M. & Wennemuth, G. Episodic rolling and transient attachments create diversity in sperm swimming behavior. BMC Biol 12, 67 (2014).

20. Leung, M.R. et al. The multi-scale architecture of mammalian sperm flagella and implications for ciliary motility. EMBO J 40, e107410 (2021).

21. Nicastro, D. et al. The molecular architecture of axonemes revealed by cryoelectron tomography. Science 313, 944–8 (2006).

22. Carbajal-Gonzalez, B.I. et al. Conserved structural motifs in the central pair complex of eukaryotic flagella. Cytoskeleton (Hoboken) 70, 101–120 (2013).

23. Gui, M. et al. De novo identification of mammalian ciliary motility proteins using cryo-EM. Cell 184, 5791–5806 e19 (2021).

24. Ichikawa, M. et al. Subnanometre-resolution structure of the doublet microtubule reveals new classes of microtubule-associated proteins. Nat Commun 8, 15035 (2017).

25. Song, K. et al. In situ structure determination at nanometer resolution using TYGRESS. Nat Methods 17, 201–208 (2020).

26. Lin, J. et al. Cryo-electron tomography reveals ciliary defects underlying human RSPH1 primary ciliary dyskinesia. Nat Commun 5, 5727 (2014).

27. Gadadhar, S. et al. Tubulin glycylation controls axonemal dynein activity, flagellar beat, and male fertility. Science 371(2021).

28. Linck, R.W., Chemes, H. & Albertini, D.F. The axoneme: the propulsive engine of spermatozoa and cilia and associated ciliopathies leading to infertility. J Assist Reprod Genet 33, 141–56 (2016).

29. Lindemann, C.B. & Lesich, K.A. Functional anatomy of the mammalian sperm flagellum. Cytoskeleton (Hoboken) 73, 652–669 (2016).

30. Warner, F.D. & Satir, P. The structural basis of ciliary bend formation. Radial spoke positional changes accompanying microtubule sliding. J Cell Biol 63, 35–63 (1974).

31. Bui, K.H., Sakakibara, H., Movassagh, T., Oiwa, K. & Ishikawa, T. Asymmetry of inner dynein arms and inter-doublet links in Chlamydomonas flagella. J Cell Biol 186, 437–46 (2009).

32. Lindemann, C.B. Functional significance of the outer dense fibers of mammalian sperm examined by computer simulations with the geometric clutch model. Cell Motil Cytoskeleton 34, 258–70 (1996).

33. Kikkawa, M., Ishikawa, T., Nakata, T., Wakabayashi, T. & Hirokawa, N. Direct visualization of the microtubule lattice seam both in vitro and in vivo. J Cell Biol 127, 1965–71 (1994).

34. Zhang, R., Alushin, G.M., Brown, A. & Nogales, E. Mechanistic Origin of Microtubule Dynamic Instability and Its Modulation by EB Proteins. Cell 162, 849–59 (2015).

35. Mitchell, D.R. & Nakatsugawa, M. Bend propagation drives central pair rotation in Chlamydomonas reinhardtii flagella. J Cell Biol 166, 709–15 (2004).

36. Suarez, S.S. Mammalian sperm interactions with the female reproductive tract. Cell Tissue Res 363, 185–194 (2016).

