## Supplementary material for "*In situ* cryo-electron tomography reveals the asymmetric architecture of mammalian sperm axonemes": Materials and Methods

All chemicals were purchased from Sigma-Aldrich unless otherwise noted.

### Sample preparation

Mouse sperms were collected from 10–15-week-old B6 wild-type mice base on the published protocol[1]. Briefly, the sperms were stripped from vasa deferentia by applying pressure to caudae epididymides in 1 x Krebs buffer (1.2 mM $KH_2PO_4$, 120 mM NaCl, 1.2 mM $MgSO_4 \bullet 7H_2O$, 14 mM dextrose, 1.2 mM $CaCl_2 \bullet 2H_2O$, 5 mM KCl, 25 mM $NaHCO_3$). The sperms were washed and resuspended in ~100 µL Krebs buffer for the following experiments.

Human sperm cells were collected by masturbation from healthy donors and visually inspected for normal morphology and motility before use. Spermatozoa were isolated by the swim-up procedure in HTF or HS solution as previously described[2] and then concentrated by 5-minute centrifugation at ≤500g and supernatant removal.

All experimental procedures utilizing human derived samples were approved by the Committee on Human Research at the University of California, Berkeley, IRB protocol number 2013-06-5395.

### Grid preparation

EM grids (Quantifoil R 2/2 Au 200 mesh) were glow discharged to be hydrophilic using an easiGlow system (Pelco). The grid was then loaded onto a Leica GP cryo plunger (preequilibrated to 95% relative humidity at 25 °C). The sperm solution was diluted and mixed with 10-nm gold beads (Electron Microscopy Science, cat #25487). Then 3.5 µL sperm mixture was added to each grid, followed by an incubation of 15 sec. The grids were then blotted for 4 sec and plunge-frozen in liquid ethane.

### Cryo-focused ion beam milling

Cryo-focused ion beam milling was performed either manually using an Aquilos cryo-FIB/SEM microscope (Thermo Fisher Scientific) or automatically using an Aquilos II cryo-

FIB/SEM microscope (Thermo Fisher Scientific). A panorama SEM map of the whole grid was first taken at 377x magnification using an acceleration voltage of 5 kV with a beam current of 13 pA, and a dwell time of 1 µs. Targets with appropriate thickness for milling were picked on the map. A platinum layer (~10 nm) was sputter coated and a gas injection system (GIS) was used to deposit the precursor compound trimethyl(methylcyclopentadienyl) platinum (IV). The stage was tilted to 15-20°, corresponding to a milling angle of 8-13° relative to the plane of grids. FIB milling was performed using stepwise decreasing current as the lamellae became thinner (1.0 nA to 30 pA, final thickness: ~300 nm). The grids were then stored in liquid nitrogen before imaging.

**Image acquisition and data processing**

Cryo-TEM tomography data of mouse sperm were collected on a 300-kV Titan Krios electron microscope (Thermo Fisher Scientific) equipped with a high brightness field emission gun (xFEG), a spherical aberration corrector, a Bioquantum energy filter (Gatan), and a K3 Summit detector (Gatan). The images were recorded at a nominal magnification of 19,500x in super-resolution counting mode. After binning over 2 x 2 pixels, the calibrated pixel size was 3.53 Å on the specimen level. For each tilt series, images were acquired using a modified dose-symmetric scheme between -48° to 48° relative to the lamella with 3° incremental steps and grouping of two images on either side (0°, 3°, 6°, -3°, -6°, 9°, 12°, -9°, -12°, 15°…). At each tilt angle, the image was recorded as movies divided into eight subframes. The total electron dose applied to a tilt series was 100 $e^{-}$/Å$^{2}$. The defocus target was set to be 4~7 µm.

Cryo-TEM tomography data of human sperm were recorded at a nominal magnification of 33,000x in super-resolution counting mode. After binning over 2 x 2 pixels, the calibrated pixel size was 2.66 Å on the specimen level. Images from a tilt series were recorded with a total dose of ~100 $e^{-}$/ Å$^{2}$. For each tilt series, images were acquired using a bidirectional scheme between -48° to 48° relative to the lamella starting from either 0° or 21°, with an incremental step of 3°. The defocus target was set to be 2~5 µm. At each tilt, the image was recorded as movies divided into eight subframes.

**Tomogram reconstruction and subvolume averaging**

All movie frames were corrected with a gain reference collected in the same EM session. Movement between frames was corrected using MotionCor2 with dose weighting[3]. Alignment of the tilt series and tomographic reconstructions were performed using Etomo[4]. The aligned tilt series were then CTF-corrected using TOMOCTF[5] and the tomograms (bin2, pixel size: 7.06 Å) were generated using TOMO3D[6]. Subsequent subvolume extraction, classification and refinement were all performed using RELION[7].

Briefly, subvolumes from the doublets were manually picked every 24 nm and extracted with a box size large enough to accommodate a complete 96-nm repeating unit (pixel size: 7.06 Å, box size: 180 pixels, dimension: 127.08 nm). All subvolumes were aligned to a map of Sea urchin doublets (EMD: 9023) lowpass filtered to 80 Å and the resulting map was used as the reference for further processing. Manual curation of the data was performed to check the alignment accuracy and data quality for each tomogram. Classification on radial spokes gave rise to 96-nm repeating units at four different registries. The doublet subvolumes were re-picked from all four class averages recentered at the base of radial spoke 2. All subvolumes were combined and aligned again to one reference and duplicate subvolumes were deleted based on distance (< 40 nm). The remaining subvolumes were aligned to yield the consensus average for all doublets. These subvolumes were later sorted and used to calculate an average of individual doublet microtubules.

To generate an average for the central pair complex, subvolumes were picked and extracted from the central pair complex every 16 nm (pixel size: 7.06 Å, box size: 160 pixels, dimension: 112.96 nm). Refinement of all subvolumes to an average of Sea Urchin central pair (EMD: 9385) resulted in a central pair with only 16-nm repeating features. The alignment parameters were modified to reset all translations (x, y, z) to zero and then used for a second round of refinement with local search only. Focused classification was performed on the microtubule-associated proteins on the C1 microtubule to separate the two populations of subvolumes (~50% each) corresponding to the central pair complex with 32-nm periodicity but separated by 16 nm. The subvolumes were then re-centered and extracted on the same protein features for the two class averages. All subvolumes were aligned to the same reference using local search. Duplicate subvolumes were deleted based on minimum separating distance (< 30 nm). The remaining subvolumes were refined to generate the consensus average for the central pair complex.

Subvolumes large enough to include all nine doublets were re-extracted based on the final positions of the subvolumes of the central pair complex (pixel size: 21.18 Å, box size: 160 pixels, dimension: 338.88 nm). These subvolumes were averaged without further alignment and used to re-extract subvolumes corresponding to individual doublets. Subvolumes corresponding to a particular doublet were re-mapped back to the tomograms. The subvolumes for 96-nm repeats obtained previously were matched and categorized into individual doublets. The manually curated subvolumes corresponding to a particular doublet were then combined and aligned.

The subvolumes corresponding to specific radial spoke-central pair interface were re-extracted based on the subvolumes of individual doublet microtubules. The radial spokes were then aligned again with a mask. This mask, plus a mask covering the central pair region, were used for the multibody analysis implemented in RELION3.

The resolutions for maps were estimated based FSC of two independently refined half datasets (FSC = 0.143). UCSF chimera was used for visualization and fitting of maps[8].

**References**


1. van der Spoel, A.C. et al. Reversible infertility in male mice after oral administration of alkylated imino sugars: a nonhormonal approach to male contraception. *Proc Natl Acad Sci U S A* **99**, 17173-8 (2002).
2. Skinner, W.M., Mannowetz, N., Lishko, P.V. & Roan, N.R. Single-cell Motility Analysis of Tethered Human Spermatozoa. *Bio Protoc* **9**(2019).
3. Zheng, S.Q. et al. MotionCor2: anisotropic correction of beam-induced motion for improved cryo-electron microscopy. *Nat Methods* **14**, 331-332 (2017).

4. Kremer, J.R., Mastronarde, D.N. & McIntosh, J.R. Computer visualization of three-dimensional image data using IMOD. *J Struct Biol* **116**, 71-6 (1996).
5. Fernandez, J.J., Li, S. & Crowther, R.A. CTF determination and correction in electron cryotomography. *Ultramicroscopy* **106**, 587-96 (2006).
6. Agulleiro, J.I. & Fernandez, J.J. Tomo3D 2.0--exploitation of advanced vector extensions (AVX) for 3D reconstruction. *J Struct Biol* **189**, 147-52 (2015).
7. Bharat, T.A. & Scheres, S.H. Resolving macromolecular structures from electron cryo-tomography data using subtomogram averaging in RELION. *Nat Protoc* **11**, 2054-65 (2016).
8. Pettersen, E.F. et al. UCSF Chimera--a visualization system for exploratory research and analysis. *J Comput Chem* **25**, 1605-12 (2004).
